# Designing optimal perturbation inputs for system identification in neuroscience

**DOI:** 10.1101/2025.03.02.641008

**Authors:** Mikito Ogino, Daiki Sekizawa, Jun Kitazono, Masafumi Oizumi

## Abstract

Investigating the dynamics of neural networks, which are governed by connectivity between neurons, is a fundamental challenge in neuroscience. Because passive (spontaneous) activity provides only limited information for estimating connectivity, perturbation-based approaches are widely applied in neuroscience, as they can evoke underlying hidden dynamics. However, the characteristics of such perturbations have typically been designed based on empirical or biological intuition. To enable more accurate estimation of connectivity, we propose a data-driven and theoretically grounded framework for optimally designing perturbation inputs, based on formulating the neural model as a control system. The core theoretical insight underlying our approach is that neural signals observed in the passive state lack sufficient latent information, which leads to failures in the system identification. Perturbations reveal these hidden dynamics and lead to improved estimation. Guided by these insights, we derive a theoretical basis for optimizing perturbation inputs that minimize estimation errors in neural system identification. Building upon this, we further explore the relationship of this theory with stimulation patterns commonly used in neuroscience, such as frequency, impulse, and step inputs. We demonstrate the effectiveness of this framework for neuroscience through simulations grounded in experimental paradigms such as neural state classification and optimal control of neural states. Our theoretical analysis, together with multiple simulations, consistently shows that perturbations designed according to our framework achieve substantially more accurate system identification compared to the conventional, intuition-based inputs. This study provides a theoretical foundation for designing perturbation inputs to achieve accurate estimation of neural dynamics. This, in turn, enables reliable discrimination of neural states such as levels of consciousness and pathological conditions, and facilitates precise control of their transitions toward recovery from abnormal states.

## Introduction

Much recent interest has focused on how interactions between individual neurons and neuronal populations support cognitive functions and behavior, and on how the disruption of these interactions contributes to various neurological and psychiatric disorders [1, 2, 3, 4, 5]. This interaction—called neuronal connectivity [6, 7]—is often represented as a network structure or a connectivity matrix. In particular, a commonly studied aspect is functional connectivity, which captures the temporal dependencies between neural activities [8, 9, 10, 11]. The functional connectivity is estimated through recording techniques such as neural spike recording techniques [12, 13], electrocorticography (ECoG) [14, 15], functional magnetic resonance imaging (fMRI) [16, 17] and electroencephalography (EEG) [18, 19]. While various methods have been proposed and used to estimate these connections (see for example [2, 20] for a comprehensive review), one typical method is based on dynamic modeling, among which is a simple but widely used linear auto-regressive model (Fig. 1a) [21, 22, 23, 24, 25]. Connectivity is statistically estimated from the time-series data of neural activity (Fig. 1b) by fitting the model parameters (Fig. 1c). A neural connectivity matrix helps visualize the connections between different brain regions, and provides a detailed map of functional interactions within the brain. By comparing connectivity matrices across individuals or groups, researchers can deepen their understanding of neuronal architectures and communication across various disciplines [8, 26, 27, 28, 29, 30].

**Fig. 1:**
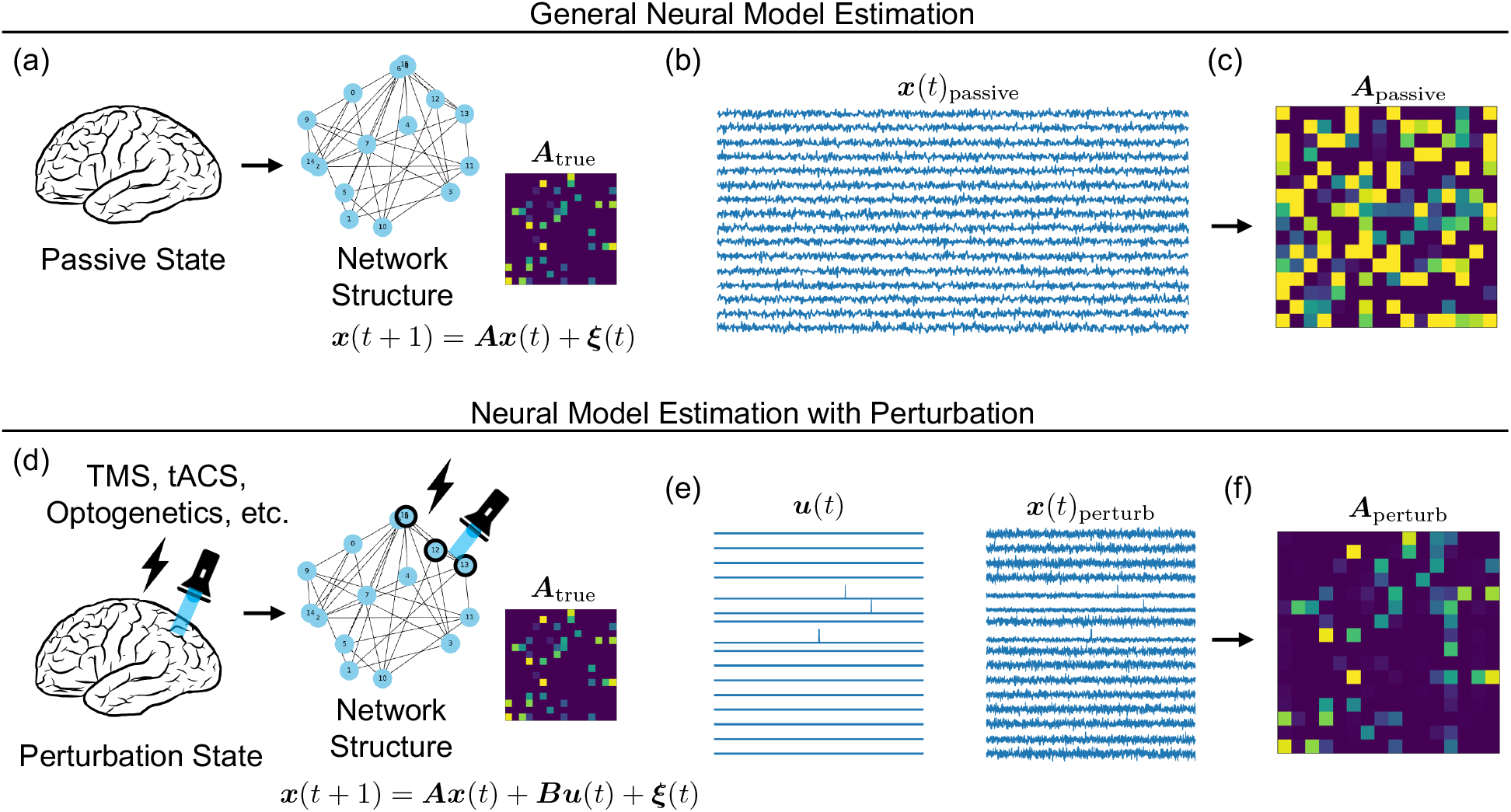
Passive and perturbation states and the resulting neural dynamics models. (a) Passive state of the neural network and the corresponding ground-truth model. (b) Recorded neural activities without perturbation. (c) Estimated model obtained without perturbation. (d) Perturbation state of the neural network and corresponding ground-truth model. (e) Recorded neural activity with perturbation. (f) Estimated model obtained with perturbation

A fundamental problem with passive observation is that it fails to reveal hidden dynamics, which leads to an invalid estimate of the corresponding parts of the model. This limitation stems from the attenuation of latent dynamical modes, such as transient or damped components. When neural activity is modeled as a linear dynamical system, as shown in Fig. 1a, the true connectivity matrix governs the generation of time-series signals (Fig. 1b). Passive observation records these spontaneous neural activities within an arbitrarily predefined temporal window and attempts to estimate the connectivity matrix using parameter estimation techniques. However, the resulting matrix ***A***_passive_ deviates significantly from the true model (Fig. 1c), because some modes quickly decay and vanish from the observable data (see also Fig. 2 for an intuitive explanation). Consequently, relying solely on passive recordings leads to misinterpretations of the neural system—such as overlooking fast transient dynamics or underestimating connectivity strength—which, in turn, may result in inaccurate conclusions in both basic neuroscience and clinical contexts.

**Fig. 2:**
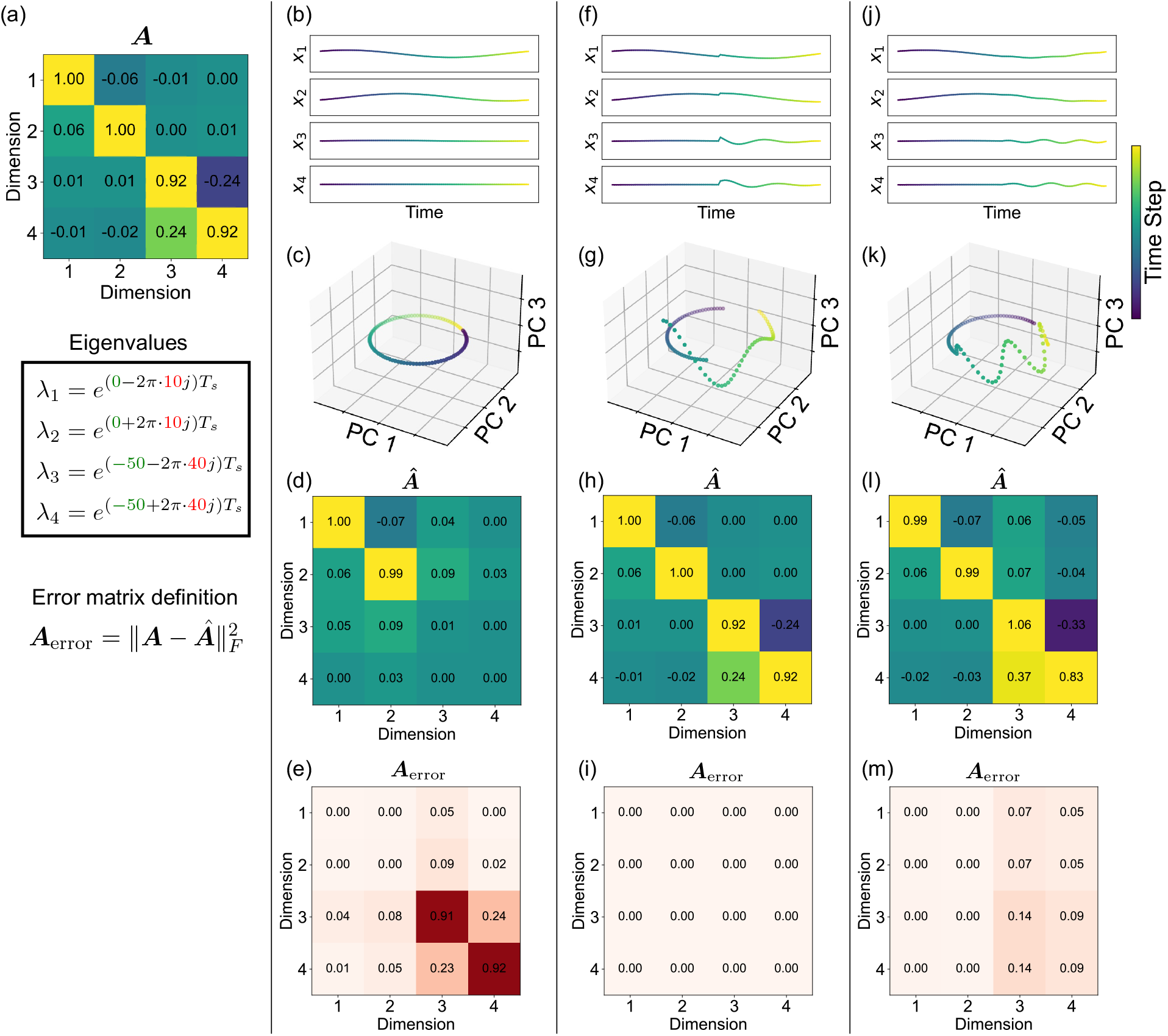
Visualization of continuous-time system dynamics by perturbations. (a) System matrix ***A*** representing the dynamics and its eigenvalues. (b) Temporal changes in the state variables of the passive state without perturbation. (c) Time-varying trajectories of each PCA component. The color gradient indicates temporal progression. (d) Estimated connectivity matrix in the passive condition. (e) Error matrix for the passive-state estimation. (f-i) Temporal changes and estimation results in response to an impulse input. (j-m) Temporal changes and estimation results in response to a sinusoidal input.

To overcome these limitations, the application of perturbations has emerged as a powerful approach for improving the estimation of connectivity in neuroscience [31, 32, 29, 33]. Its effectiveness lies in its ability to actively manipulate the neural system, rather than merely observing it, and to thereby uncover dynamic and causal interactions that are otherwise hidden [32]. Recent studies using stimulation techniques such as optogenetics [34, 35] and transcranial magnetic stimulation (TMS) [36, 29, 37] have demonstrated that these interventions can significantly improve the detection of causal relationships within the neural system. Such approaches have been applied to the study of cognitive processes and states of consciousness [31, 29]. These studies underscore the value of perturbation inputs in driving state transitions in the neural system, thereby enabling more accurate and reliable connectivity estimation.

However, the parameters of perturbation protocols, such as the location, intensity, and shape of stimulation, have often been determined based on anatomical and physiological insights and on empirically established techniques [38, 39, 29, 40, 41]. While these approaches have provided practical utility, they may not optimally exploit the underlying neural dynamics or maximize the informativeness of the perturbation. To overcome these limitations and enhance the reliability of connectivity inference, there is a pressing need to design data-driven and theoretically grounded perturbations.

In this paper, we propose a framework for designing the optimal perturbation input through control theory in neuroscience. We interpret neural dynamics as a control system [42, 43, 44, 45, 46, 47], and treat external perturbations as control inputs to design properties of neural stimulation (Fig. 1d). If the optimal perturbation input can be systematically designed, it becomes possible to steer the neural system toward states that are maximally informative (Fig. 1e), thereby enhancing the accuracy of the inferred connectivity (Fig. 1f). While recent studies have begun to develop algorithmic approaches to active stimulation design for specific experimental platforms [48, 49], a general theoretical framework for why and which perturbation inputs are effective has yet to be established. We first describe how to formulate neural dynamics as a control system and how to estimate the model parameters from observed data. Building upon this formulation, we derive a theoretical basis that enables us to design the optimal perturbation inputs for the neural system identification. We demonstrate the validity and utility of this theoretical basis by exploring its implications for optimizing parameters of common neurostimulation techniques and by applying it to practical examples, including neural state classification [31, 36, 50, 47] and control of neural states [42, 51, 44, 45]. In these demonstrations, we define concrete problems and apply the theory to validate its practical utility.

Our research offers a comprehensive framework for system identification with perturbation in neuroscience, and paves the way for more precise and effective analysis of neural dynamics. By providing clear guidelines on the design and application of perturbations, this framework serves as a practical reference for experimental settings, helping researchers determine the most effective stimulation parameters for their studies.

## Background

To evaluate how external perturbations enhance the accuracy of system identification, this section reviews the estimation formulations for linear dynamical systems, comparing the cases with and without perturbation inputs.

### System Identification with External Perturbation

This section reviews a well-established framework for system identification of linear dynamical systems with external perturbations. Researchers have modeled brain dynamics as a discrete-time linear dynamics with external inputs and stochastic system noise [52, 42, 53, 54, 43]. We assume that the brain state evolves according to the following linear dynamics:

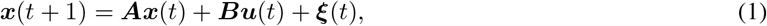

where ***x***(*t*) ∈ ℝ^*n*^ denotes the neural state vector at time *t*, such as the activity of neurons, populations, or brain regions, and *n* represents its dimension. The connectivity matrix ***A* ∈** ℝ^*n×n*^ characterizes intrinsic interactions reflecting functional connectivity. The input matrix ***B* ∈** ℝ^*n×m*^ specifies how external inputs ***u***(*t*) **∈** ℝ^*m*^—including sensory stimuli or interventions such as TMS—affect the system, where *m* represents the number of perturbation channels. The input ***u***(*t*) is freely designed and known to the experimenter. Finally, ***ξ***(*t*) ℝ^*n*^ represents stochastic fluctuations (e.g., synaptic noise or unobserved inputs), modeled as i.i.d. zero-mean Gaussian noise across time with covariance matrix **Σ**_*ξ*_ = 𝔼 [***ξ***(*t*)***ξ***(*t*)^⊤^] ≻ 0. We remark that neural activities measured at the macroscopic level, such as functional magnetic resonance imaging (fMRI) or intracranial electroencephalography (iEEG) have been experimentally and theoretically validated to follow approximately linear dynamical systems in Refs. [55, 56].

Here, we assume the input matrix ***B*** is known, and we are only estimating the connectivity matrix ***A*** from the time series data of ***x*** and ***u***. We construct the data matrices as

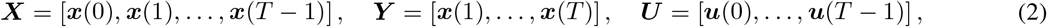

where *T* represents the length of the time series data. Using these matrices, the parameter matrix ***A*** can be estimated by minimizing the reconstruction error in the sense of ordinary least squares (OLS):

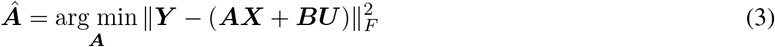

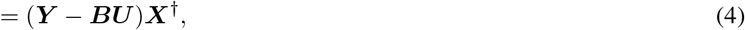

where ∥ · ∥_*F*_ represents the Frobenius norm and the symbol ^*†*^ denotes the pseudoinverse of a matrix. This manuscript derives a theoretical framework based on this formulation, which assumes linear dynamics with the input matrix ***B***— known. The relaxation of these assumptions—nonlinear state dynamics and the case of unknown ***B***—is addressed in Supplementary Material B.2 and B.3.

### System Identification without External Perturbation

To compare the quality of system identification with and without perturbations, we also consider the passive case without external input. This corresponds to setting ***u***(*t*) = **0** in the formulation in the perturbed case, yielding the following dynamics:

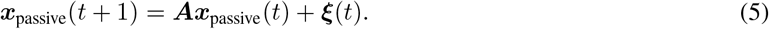

Here, ***A*** and ***ξ***(*t*) are identical to those defined in Eq. 1. Because this system evolves differently from the perturbed case, we denote its state trajectory as ***x***_passive_(*t*) to explicitly distinguish it from the dynamics under perturbation.

Similarly, the parameter matrix ***A*** in the passive condition can be estimated by applying the ordinary least squares (OLS) method:

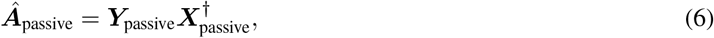

where

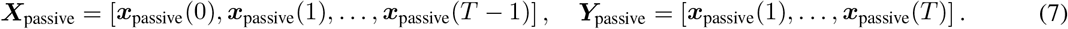

The subscript “passive” is used again to explicitly distinguish variables associated with the unperturbed dynamics from those obtained under external perturbations.

### Neural Stimulation as Control Inputs

This section describes how commonly used neural stimulation techniques can be related to input signals in control theory. Their adjustable parameters vary depending on how the stimulation inputs are modulated.

Three non-invasive electrical stimulation methods illustrate how stimulation paradigms map onto basic control inputs. Transcranial magnetic stimulation (TMS) induces brief and transient perturbations via electromagnetic pulses [57], which are naturally represented as a sequence of impulse-like inputs, where the timing and intensity of each pulse are the primary controllable parameters. Transcranial direct current stimulation (tDCS) primarily modulates neural activity through approximately constant inputs [58], which can be viewed as a step-like signal whose main controllable parameter is the amplitude of the applied current. Transcranial alternating current stimulation (tACS) delivers oscillatory inputs [59], corresponding to sinusoidal signals characterized by amplitude, frequency, and phase. In control theory, impulse, step, and sinusoidal inputs are the basic components used to characterize system responses and dynamics [60, 61].

The control input framework extends beyond non-invasive techniques to invasive and optogenetic stimulation. Invasive electrical stimulation, including intracranial microstimulation and deep brain stimulation (DBS), enables direct delivery of electrical inputs to neural tissue [62], providing flexible control over amplitude and timing through pulse trains or temporally structured waveforms. Optogenetic stimulation allows genetically targeted activation or inhibition of specific neurons using light [63], providing fine-grained control over multiple input dimensions, including amplitude (light intensity), temporal pattern, and cell-type specificity. In particular, recent developments enable stimulation at the level of individual neurons with high temporal precision [64, 65], allowing flexible construction of spatiotemporal input patterns.

These stimulation examples demonstrate that the theoretical framework developed in this paper connects to practical experimental settings. While a substantial gap remains between idealized control inputs in theory and experimentally realizable stimulation, the core principles established in the following sections provide a foundation that naturally extends to these practical stimulation paradigms.

## Results

We present a theoretical framework for determining the optimal perturbation inputs to enhance neural system identification. We begin by presenting an intuitive example in which the failure of passive estimation is demonstrated, and the benefit of perturbation inputs in re-exciting weakly observable dynamics becomes evident. Following this, we provide a guiding theoretical principle that estimation error of ***A*** is reduced by excitation of the state ***x*** by perturbation input ***u***. We then validate this theoretical foundation by applying it to canonical perturbation signals used in neuroscience—sinusoidal inputs and impulse as typically observed in tACS, tDCS, and TMS—and derive analytical relationships between input parameters and estimation errors. Furthermore, we demonstrate the applicability of the proposed framework to practical problems in neuroscience, such as neural state classification and optimal control for neural state transitions. Finally, we outline a practical framework for designing optimal perturbation inputs to estimate neural dynamics.

### Typical Failure of Passive-State Model Estimation

We demonstrate why passive state observation, a common experimental condition in system neuroscience, fails to estimate the system model of neural dynamics. Although passive observation is widely employed due to its convenience and non-invasiveness, this practice has fundamental limitations: it inevitably overlooks causal and dynamical information that vanishes under spontaneous conditions but can be recovered through external perturbations, leading to degraded connectivity estimates. Such limitations lead to misinterpretations of the underlying neural functions when neural dynamics are modeled as a control system.

We estimate the connectivity matrix from simulated time series data generated by the true model. The true model, characterized by a connectivity matrix ***A***, is illustrated in Fig. 2a. Neural data are generated based on the connectivity matrix, yielding the time series shown in Fig. 2b. These data are not influenced by external input (passive state), and thus the oscillations of *x*_3_ and *x*_4_ gradually attenuate over time. This attenuation is further confirmed by principal component analysis (PCA), which reveals that the first and second principal components capture the 10Hz dynamics, while the third principal component, associated with the 40Hz mode, contributes negligibly to the total variance (Fig. 2c). Consequently, the matrix estimated by OLS deviates substantially from the true connectivity matrix (Fig. 2d), particularly showing errors in the lower-right components, as highlighted in Fig. 2e. This failure arises because the true matrix ***A*** contains two distinct dynamical modes: one is a continuous oscillatory mode at 10Hz, and the other is a damped mode at 40Hz. The damped mode becomes unobservable in the time series once its contribution has attenuated and vanished. Passive recordings capture only superficial aspects of the connectivity and fail to estimate the true underlying neural dynamics.

The limitation of the passive state can be resolved by applying perturbation inputs. Figures 2f–2i show the application of an impulse input ***u*** to the system. This perturbation input primarily affects the nodes corresponding to *x*_3_ and *x*_4_, reintroducing variations in the third principal component direction. Estimating matrix ***A*** from this perturbed time-series data yields a more accurate estimate than in the passive case. Similarly, an improvement is observed when different types of input—a sinusoidal input for example—are used instead of the impulse input, as shown in Figs. 2j–2m. These results illustrate that appropriate perturbation inputs can effectively re-excite weak dynamics, thereby enhancing system identification accuracy. This can be theoretically supported by the concept of persistent excitation [66, 67, 68, 69]. According to this theory, a perturbation input is said to be persistently exciting if it causes the system’s state to sufficiently explore the state space over time. These insights underscore the importance of incorporating controlled stimulation in experimental design, particularly when accurate system identification is desired. A similar trend under nonlinear dynamics is shown in Supplementary Material B.2.

### Intuitive Perturbation Design Informed by Covariance Eigenvalues

In this section, we show how perturbations reduce the estimation error of the system matrix ***A*** by exciting the state dynamics, thereby providing a theoretical basis for designing effective perturbation inputs. When the data length *T* is sufficiently large (*T* → ∞), the estimation error of ***A*** asymptotically converges to the following expression [70]:

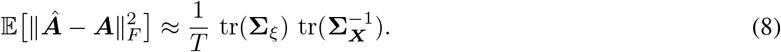

where 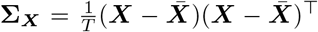 with 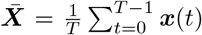 represents the covariance matrix of the state vector ***x***, and tr denotes the trace of a matrix. The detailed derivation of this equation is provided in the Supplementary Material A.1. This equation indicates that the estimation error of ***A*** can be reduced either by increasing the measurement duration *T* or by minimizing the trace of the inverse covariance matrix 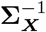, which depends on the perturbation input ***u***(*t*). Intuitively, this relationship resembles a signal-to-noise ratio, where increasing the covariance of the state dynamics relative to the noise covariance leads to improved estimation accuracy.

The asymptotic expression in Eq. 8 reveals that the estimation error of ***A*** depends inversely on the eigenvalues of the covariance matrix **Σ**_***X***_. Expressing this relationship explicitly in terms of the eigenvalues yields

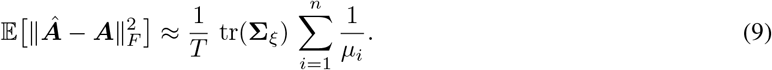

where *µ*_*i*_ denotes the *i*-th eigenvalue of **Σ**_***X***_. This relationship indicates that small eigenvalues dominate the estimation error through their inverse contributions. Therefore, the perturbation input should be designed not only to increase the overall variance of the state ***x***, but to enlarge the eigenvalues of **Σ**_***X***_ more uniformly across all directions of the state space. In other words, exciting all dynamical modes of the system–rather than amplifying a limited subset–is essential for improving estimation accuracy.

This theoretical insight can be illustrated intuitively in Fig. 3. The figure visualizes ellipsoids constructed from the eigenvalues *µ*_*i*_ and their reciprocals 1*/µ*_*i*_ of the covariance matrix **Σ**_***X***_. The covariance matrices **Σ**_***X***_ are calculated from the time series of the state vector ***x*** obtained under passive and perturbation conditions (Fig. 3a). The length of each axis of the ellipsoid corresponds to the variance of the state along that direction, which is associated with the eigenvalue *µ*_*i*_. Its reciprocal counterpart 1*/µ*_*i*_ represents the contribution to the estimation error (Fig. 3b). As shown in Fig. 3c, when the neural dynamics is dominated by a single eigenvalue *µ*_1_, the other eigenvalues *µ*_2_ and *µ*_3_ remain small, resulting in an elongated reciprocal ellipsoid (Fig. 3d) and a large estimation error. When perturbation inputs that excite the directions associated with *µ*_2_ and *µ*_3_ are applied, the ellipsoid expands and becomes closer to a sphere (Fig. 3e). As a result, the estimation error decreases in all directions, as illustrated in Fig. 3f. This uniform enlargement of the eigenvalues provides a clear guideline: perturbations should be designed so that the variance becomes large in all directions, rather than being confined to specific modes.

**Fig. 3:**
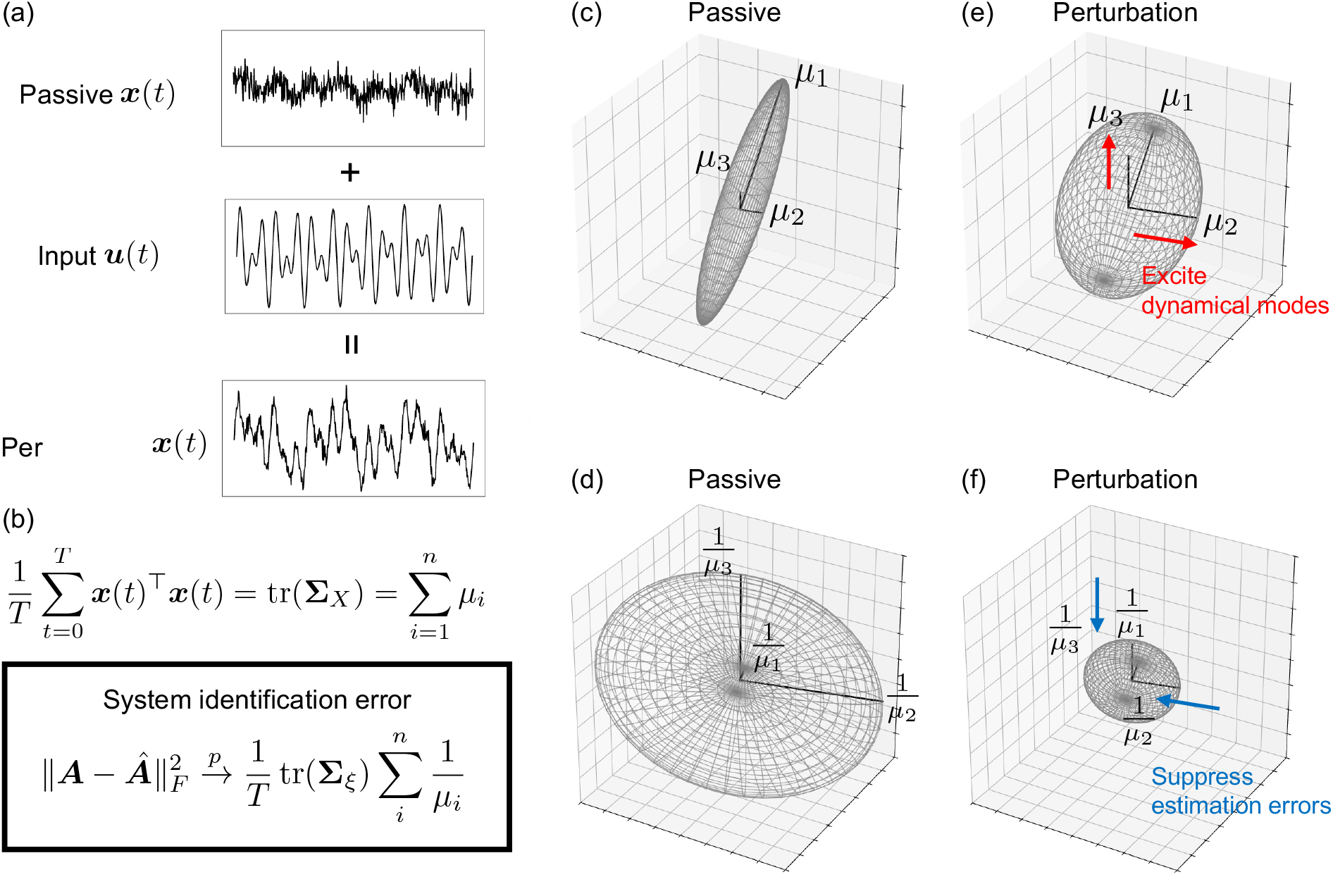
Effect of perturbation on the eigenvalues of the state covariance matrix. Ellipsoidal plots show the eigenvalues and reciprocal eigenvalues of the covariance matrix of the state trajectories ***X***, comparing passive dynamics and the case with perturbation. All ellipsoidal plots (c–f) are drawn along the first three eigenvector directions of **Σ**_***X***_; panels (c, e) scale the axes by the eigenvalues *µ*_*i*_, while panels (d, f) scale them by the reciprocals 1*/µ*_*i*_. (a) Examples of a passive signal, a perturbation input, and a perturbed signal. (b) Analytical relationship linking the eigenvalues of the state covariance matrix to system identification error. (c) Passive: The covariance matrix is dominated by a single eigenvalue direction. (d) Passive (reciprocal): A single dominant eigenvalue produces a widely spread reciprocal-space ellipsoid (e) Perturbation: Additional dynamical modes are excited, increasing the smaller eigenvalues. (f) Perturbation (reciprocal): Perturbations yield a more uniform reciprocal-space ellipsoid, reducing estimation error.

### Theoretical Formulation of Sinusoidal Inputs for an Overall Increase in Eigenvalues

Based on the guideline presented in the previous section, we derive analytical expressions to investigate which frequency of perturbation input will efficiently excite the dynamical modes of a system. In neuroscience, such a perturbation is analogous to tACS. Since neural dynamics possess modal frequencies, there should exist perturbation input frequencies that resonate with these modes. Such resonance leads to an amplification of ***x***(*t*), which generally results in larger eigenvalues *µ*_*i*_ of the covariance matrix **Σ**_***X***_, reflecting increased variability along the corresponding modes. To clarify this relationship, we derive an analytical expression for ***x***(*t*) to investigate how the input frequency affects its amplitude. The solution ***x***(*t*) to the model in Eq. 1 can be obtained by iterating the state transition matrix. Specifically,

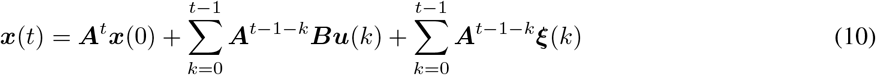

We assume a cosine input 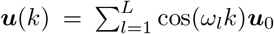, where *ω*_*l*_ are the angular frequencies. By substituting this into the second term, we can obtain the following equation by performing diagonalization of ***A*** and expressing the eigenvalues in polar form.

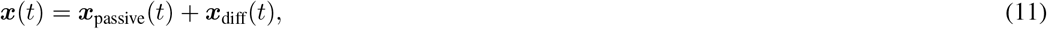

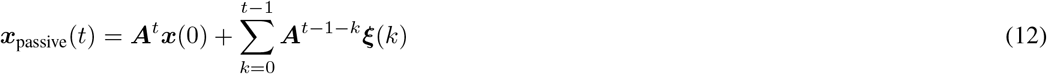

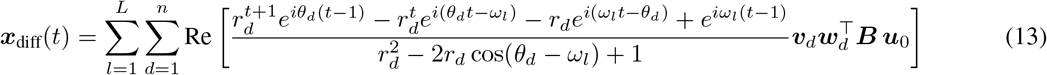

where each eigenvalue of ***A*** is written in polar form as 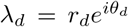, with *r*_*d*_ = *λ*_*d*_ denoting the magnitude and *θ*_*d*_ = arg(*λ*_*d*_) denoting the argument. ***x***_passive_ represents the state ***x*** that would be realized if no perturbation were applied, i.e., the state that would evolve according to Eq. 5. The vector ***v***_*d*_ denotes the right eigenvector of ***A*** associated with *λ*_*d*_, and 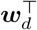 denotes the corresponding left eigenvector. The detailed derivation is provided in the Supplementary Material A.2.1.

We focus on ***x***_diff_(*t*) simply because it is the component directly driven by the control input. When the input frequencies *ω*_*l*_ coincide with the mode’s angular component *θ*_*d*_, a resonance-like amplification occurs in ***x***_diff_(*t*)—that is, the amplitude of ***x***(*t*) increases markedly. Increasing the magnitude of ***x***(*t*) through resonance naturally leads to an increase in the overall variance in **Σ**_***X***_. This enhancement directly contributes to increasing all eigenvalues *µ*_*i*_ of the covariance matrix rather than leaving some unexcited. Therefore, the analysis of Eq. 13 provides the necessary guidelines to target and amplify specific dynamical modes, suggesting that oscillatory inputs, such as tACS, should be designed with frequencies that correspond to the true dynamical mode frequencies.

### Theoretical Formulation of Impulse and Step Inputs

The eigenvalues of the covariance matrix can also be increased by manipulating the intensity of the perturbation input. In neuroscience, external perturbations such as TMS and tDCS can be modeled as impulse-like or step-like inputs to neural systems. The state vector ***x***(*t*) for impulse inputs can be written as:

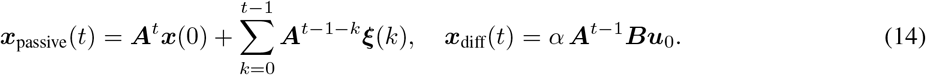

For step input, the state vector is described by,

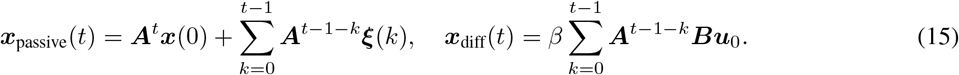

where *α* and *β* are intensity of the impulse and step inputs. The detailed derivation is provided in the Supplementary Material A.2.2. From these equations, it is clear that the contribution of the perturbation to the state covariance increases proportionally to the squares of the input strengths, *α*^2^ and *β*^2^. However, estimation accuracy is not determined by input intensity alone. Eqs.14 and 15 show that the resulting dynamics ***x***_diff_(*t*) are critically dependent on the term ***Bu***_0_. This term represents how the input vector ***u***_0_ (which defines the spatial pattern of the stimulation, i.e., which nodes are targeted) interacts with the system’s input matrix ***B***. Therefore, while increasing intensity (*α, β*) within experimental constraints is beneficial, the spatial pattern ***u***_0_ is the key design parameter that determines which dynamical modes are excited. An improperly chosen ***u***_0_ may excite only the modes already dominant in the passive state, failing to enlarge the smaller eigenvalues that are critical for system identification. The problem of how to design the optimal spatial pattern ***u***_0_ to most effectively excite the weakly observable dynamics will be addressed in a later section.

### Demonstration of Sinusoidal Inputs

In this section, we demonstrate how to design the frequency of oscillatory input by analyzing the eigenvalues of the covariance matrix. Theoretical formulations indicate that tuning the frequency of oscillatory perturbation inputs is more complex compared to that of impulse and step inputs. We generate neural dynamics governed by the connectivity matrix ***A***, which is characterized by designated dynamical modes and compare its eigenvalues *λ*_*i*_ with eigenvalues *µ*_*i*_ of the covariance matrix of the perturbed state vectors. The demonstration for impulse and step inputs can be found in the Supplementary Material B.4.

The simulation results show that an oscillatory input with the same frequency as a mode of the system matrix minimized the estimation error when the system had a single mode. Figure 4a shows the system matrix and its eigenvalues, which correspond to a single 10 Hz mode. We generated time series data using this system matrix and estimated the matrix. Figure 4b illustrates the changes in the sum of eigenvalues of the state covariance matrix **Σ**_***X***_, denoted as ∑*µ*_*i*_, the sum of reciprocal eigenvalues 1*/*∑*µ*_*i*_, and the estimation error defined by the Frobenius norm. The sum of eigenvalues is maximized by a 10 Hz sinusoidal input, while the sum of reciprocal eigenvalues is minimized. Consistent with these results, the estimation error is minimized by the 10 Hz sinusoidal input. As shown in Eq. 9, the estimation error is proportional to the sum of the inverse eigenvalues. This tendency can be explained more clearly by plotting the eigenvalues as ellipsoids. We visualized the eigenvalues using eigenvalue-scaled ellipsoids, as shown in Figs. 4c and 4d. Both the major and minor axes of the ellipsoids in Fig. 4c are extended by the sinusoidal inputs, especially by the 10 Hz input. In contrast, the major and minor axes of the ellipsoids in Fig. 4d are substantially shortened by the 10 Hz sinusoidal input.

**Fig. 4:**
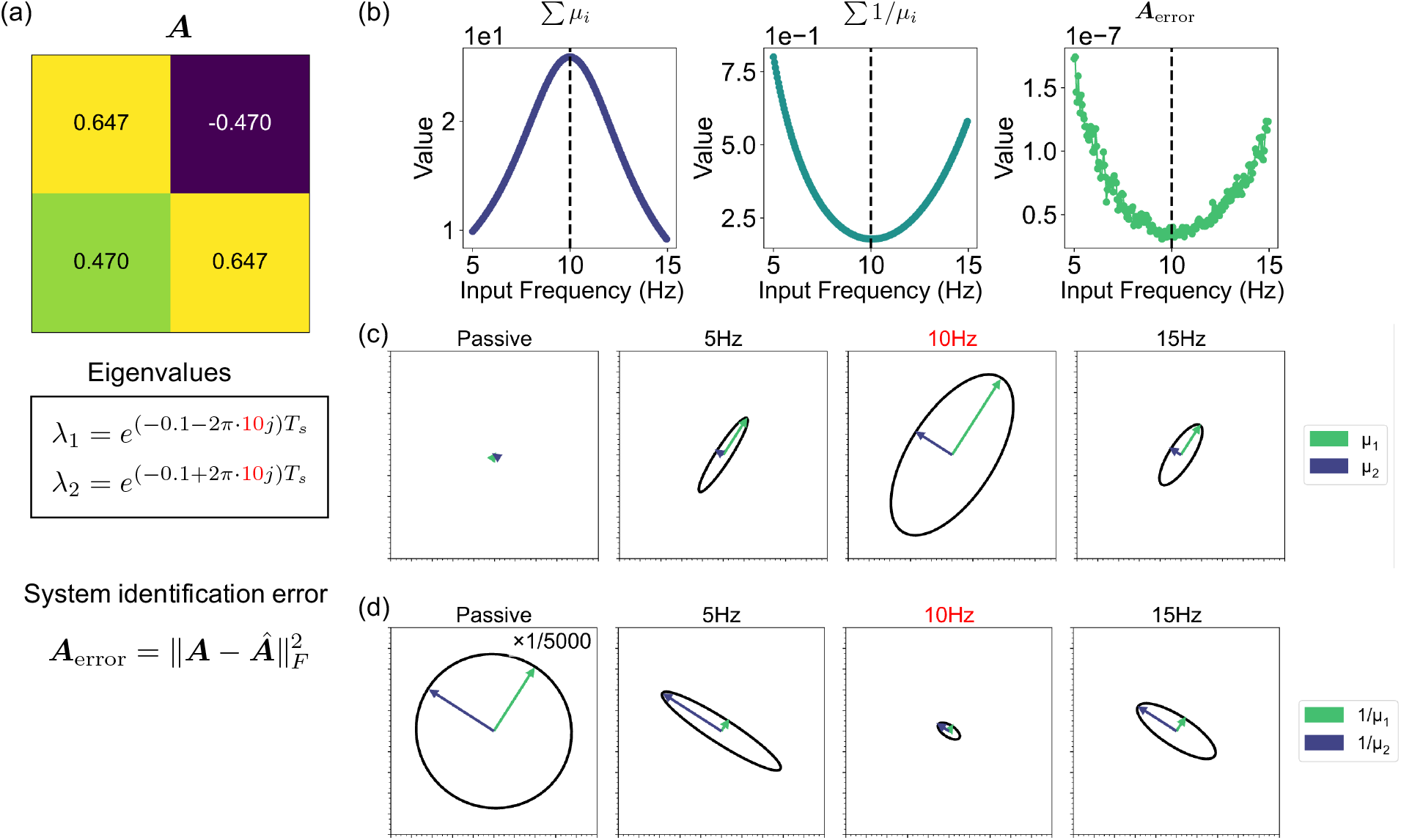
Error in the system matrix and eigenvalue-scaled ellipsoids. (a) State transition matrix ***A*** and its eigenvalues *λ*_*i*_. (b) System frequency responses, showing the sum of eigenvalues of the state covariance matrix **Σ**_***X***_ (left), the sum of inverse eigenvalues of the state covariance matrix **Σ**_***X***_ (middle), and the system matrix error (right). (c) Eigenvalue ellipsoids of **Σ**_***X***_ under passive, 5 Hz, 10 Hz, and 15 Hz conditions. (d) Reciprocal-eigenvalue ellipsoids of **Σ**_***X***_ under passive, 5 Hz, 10 Hz, and 15 Hz conditions.

When the system exhibits multiple modes, the input frequencies should be designed to reflect all these modes. We demonstrate this using the system matrix shown in Fig. 5a, along with simulated time series and the corresponding matrix estimation. Because the system matrix contains two distinct pairs of eigenvalues (10 Hz and 20 Hz), we constructed the evaluation input ***u*** as a combination of two frequencies. The sum of eigenvalue reciprocals is minimized only when the input contained both 10 Hz and 20 Hz components; inputs with a single frequency yield larger values (Fig. 5b). To further evaluate the effect of input design, we compared the two-frequency input with a flat-spectrum input (Fig. 5c). When the total input energy was normalized, the two-frequency input achieves better identification performance than the flat-spectrum input, which is commonly employed in control and system identification studies. This result highlights the importance of tailoring the input spectrum to the system’s intrinsic dynamics rather than relying on uniform excitation. The superior performance of the two-frequency combination can be understood through the eigenvalue-scaled ellipsoids in Figs. 5d–g. Figure 5d illustrates ellipsoids determined by the three dominant eigenvalues. The ellipsoid volume is expanded by three types of sinusoidal inputs (10 Hz, 20 Hz, and the combined input). However, the third axis (blue line) is not extended under single-frequency inputs, as shown in Fig. 5e, leading to larger estimation errors. Small eigenvalues disproportionately increase the sum of reciprocal eigenvalues, as seen in Figs. 5f and 5g. In these cases, the reciprocal eigenvalue-scaled ellipsoids are enlarged along the third axis (blue line) when only a single-frequency input is applied. By contrast, the combined input uniformly increases all eigenvalues, including the third, thereby successfully minimizing the reciprocal eigenvalue-scaled ellipsoid volume. These findings indicate that sinusoidal inputs should be designed to match multiple system modes rather than single modes. It should be noted that the true system modes cannot generally be known a priori. They must be identified through iterative experiments and refinement of the estimated system matrix. This practical procedure for optimal perturbation design is described in a later section (see Practical Designing Procedure for Optimal Perturbation Inputs).

**Fig. 5:**
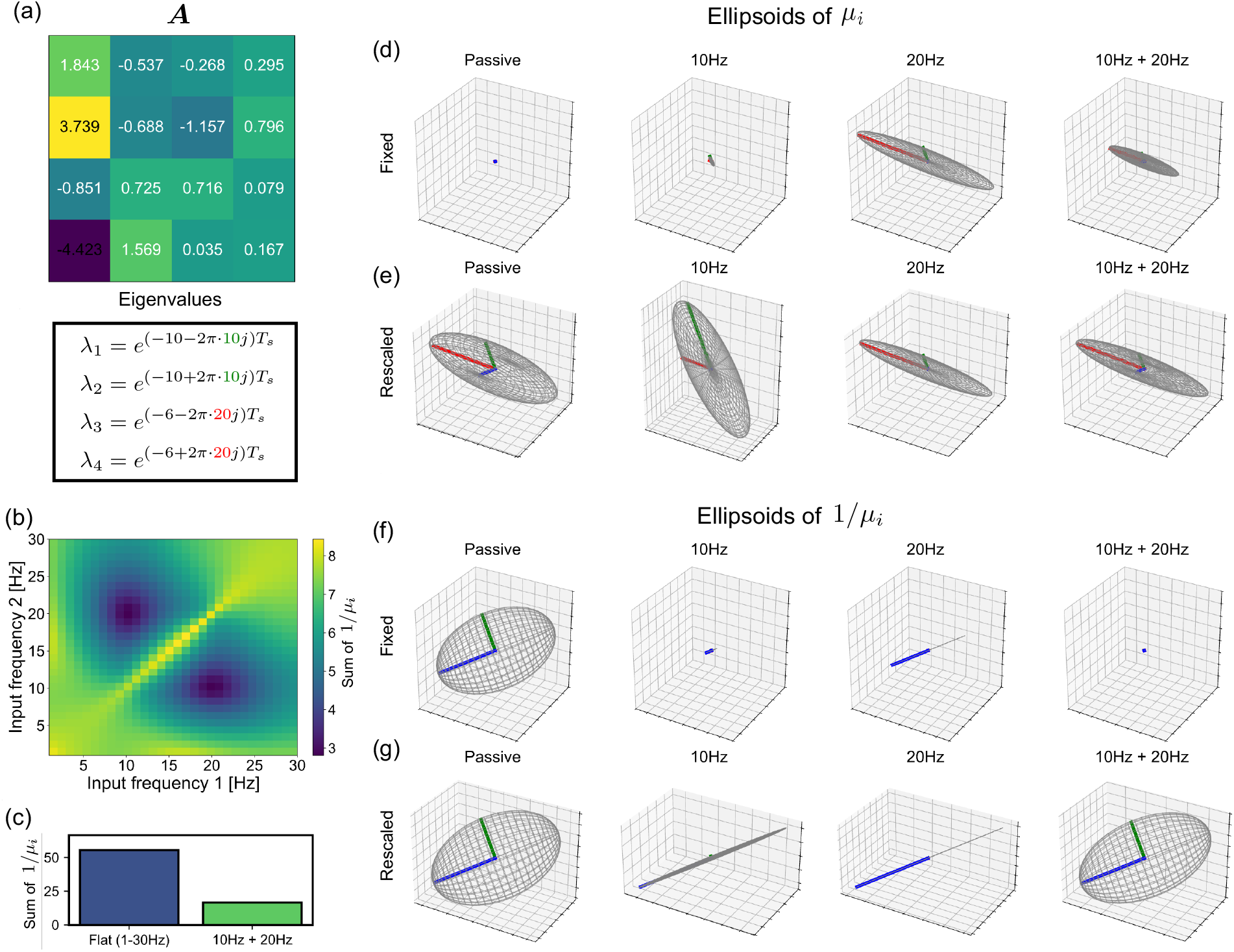
Simulation of a multi-mode system and eigenvalue-scaled ellipsoids. (a) System matrix ***A*** and its eigenvalues; the system exhibits two damped oscillatory modes at 10 Hz and 20 Hz. (b) Theoretical estimation error, defined as ∑ (1*/µ*_*i*_) for two-frequency input combinations; cooler colors indicate smaller estimation errors. (c) Comparison of the reciprocal eigenvalue sum for flat-spectrum versus composite-frequency inputs. (d) Eigenvalue-scaled ellipsoids (*µ*-scaled) under four input conditions (columns): Passive, 10 Hz, 20 Hz, and 10 Hz + 20 Hz; ellipsoid axes align with the eigenvectors. (e) Relative-scale view of (d). (f) Eigenvalue-scaled ellipsoids (inverse-scaled, 1*/µ*_*i*_) for the same four conditions. (g) Relative-scale view of (f).

### Demonstration of Location Tuning

This section illustrates how our theoretical framework enables location-specific tuning of perturbation inputs within neural networks. As established earlier, perturbations should be designed so that the variance becomes large in all directions, rather than being confined to specific modes. Two candidate strategies arise naturally: directly stimulating the nodes associated with heavily damped modes to target those specific modes, or stimulating a hub node whose outgoing connections propagate the perturbation across the entire network.

To examine this relationship in practice, we constructed a network as shown in Figs. 6a and 6b. The network consists of seven nodes, structured into three oscillatory modes (Nodes 1–6) and one hub node (Node 7). It is designed to have three oscillatory modes at 15, 25, and 35 Hz. Fig. 6c shows the damping rates |*λ*_*A*_| of the eigenvalues of ***A***: the mode formed by Nodes 1 and 2 (15 Hz) is heavily damped, while those formed by Nodes 3–6 are moderately damped. Node 7 serves as a hub with only outgoing edges. Each node was individually perturbed by an impulse input. Here, an impulse input is defined as a Kronecker delta at *t* = 0 with fixed amplitude *α* = 10, with no external input applied at any subsequent time step.

**Fig. 6:**
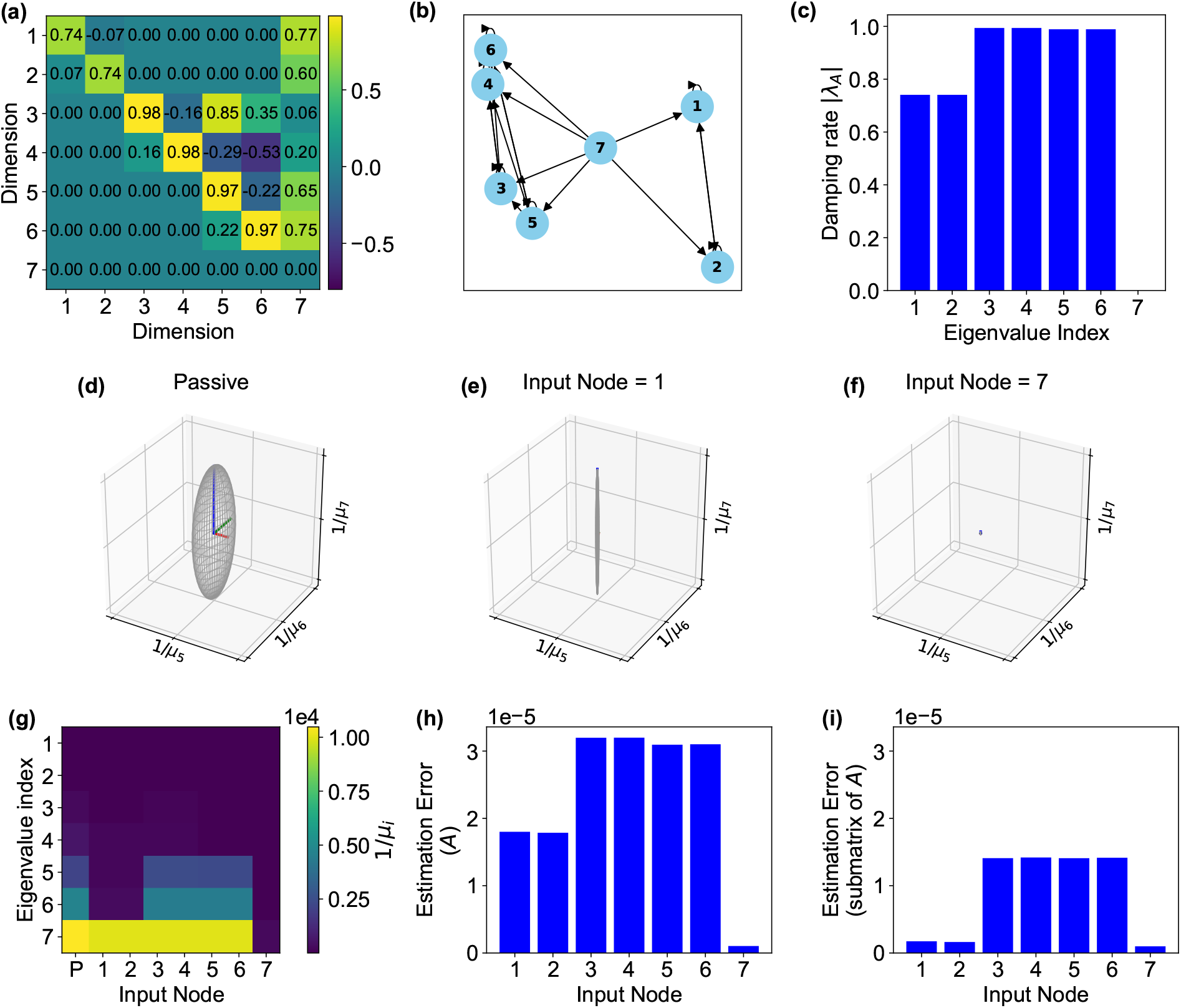
**Optimal stimulation location (a hub node or the heavily-damped subnetwork) minimizes estimation error**. A 7-node network with three oscillatory modes (15, 25, 35 Hz) is used to identify effective perturbation locations. The 15 Hz mode (Nodes 1–2) is heavily damped; Node 7 is a hub with only outgoing edges. (a) The 7 × 7 connectivity matrix ***A***. (b) Graph representation. (c) Damping rates |*λ*_*A*|_ of the eigenvalues of ***A***. (d–f) 3D ellipsoids whose axes are proportional to 1*/µ*_5_, 1*/µ*_6_, 1*/µ*_7_ (reciprocals of the three smallest eigenvalues of **Σ**_***X***_; larger axes indicate greater estimation error): (d) passive, (e) impulse to Node 1, (f) impulse to Node 7. (g) Heatmap of 1*/µ*_*i*_ across input nodes (P = passive); uniformly small values appear only for Node 7. (h) Full-matrix estimation error per input node. (i) Estimation error for the submatrix excluding Node 7; Nodes 1, 2, and 7 achieve comparable errors, confirming that direct stimulation of the damped subnetwork matches hub stimulation.

Stimulating Node 7 (hub node) minimizes the total estimation error of ***A*** (Fig. 6h), because its outgoing connections propagate the perturbation to all nodes and re-excite every dynamical mode. Figure 6d–f illustrate how the choice of input node shapes the smallest eigenvalues of **Σ**_***X***_. Because the 15 Hz mode (Nodes 1–2) is heavily damped, it decays rapidly and contributes almost no variance to **Σ**_***X***_ under passive conditions; as a result, the fifth through seventh eigenvalues *µ*_5_–*µ*_7_ are small and their reciprocals 1*/µ*_5_–1*/µ*_7_ are large, producing the elongated ellipsoid seen in Fig. 6d. When Node 1 receives an impulse (Fig. 6e), the eigenvalue associated with the 15 Hz mode grows and the 1*/µ*_5_ and 1*/µ*_6_ axes shrink; however, the 1*/µ*_7_ axis—which correspond to directions that Node 1 alone does not excite— remains large. Stimulating Node 7 (Fig. 6f) re-excites all six oscillatory nodes through its outgoing connections, simultaneously reducing all three axes. Figure 6g confirms this pattern across all candidate nodes: the contribution 1*/µ*_*i*_ remains uniformly small only when Node 7 receives the impulse. Figures 6h and 6i show that Node 7 achieves the smallest full-matrix estimation error, followed by Nodes 1 and 2, which directly drive the heavily damped 15 Hz mode.

As this simulation represents only one example of location design, its limitations and the corresponding countermeasures should be stated. In actual experiments, the most effective perturbation location depends on factors such as hub-node connectivity and modal damping rates, and ***B*** itself may not always be known a priori. In such cases, approaches such as iterative optimization of 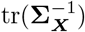 (described in a later section) and joint estimation of ***A*** and ***B*** (Supplementary Material B.3) provide systematic alternatives. Nevertheless, the results presented here provide an intuitive guideline: stimulation directed at hub nodes or at nodes driving heavily damped modes effectively excites the full set of dynamical modes and minimizes estimation error.

### Neural State Classification

We demonstrate the effectiveness of our framework by applying it to a neural state classification problem. Neural state classification is crucial for diagnosing illnesses and assessing brain conditions in many experimental neuroscience settings [31, 36, 50, 47]. Some of these studies have already applied arbitrary perturbation inputs for classification, but not optimal perturbation inputs. By simulating such real-world applications, we demonstrate how well optimal perturbation design can contribute to neuroscience research.

We designed a neural network with clearly distinct task conditions and considered a simulation setting in which these conditions are classified using signals of a fixed duration. These distinct task conditions consist of five types, each defined by a unique linear dynamical system characterized by differing eigenvalue spectra and connectivity topologies of matrix ***A*** (Fig. 7a). These task conditions are intended to mimic different cognitive or behavioral contexts. For example, in a typical motor task experiment, such conditions could correspond to motor execution or imagery involving the left or right hand, or resting state [50, 47]. The neural signals were simulated under five different task conditions and two stimulation conditions: passive observation and external perturbation. Perturbation was applied as impulse-type inputs, such as TMS. The stimulus location was determined for each task condition by applying an impulse to each node and selecting the one that minimized 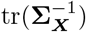. The resulting time-series data are shown in Fig. 7b. Using this data, we estimated the underlying dynamical system via a control-based identification approach presented in Eq. 4, which corresponds to an estimation of functional connectivity.

**Fig. 7:**
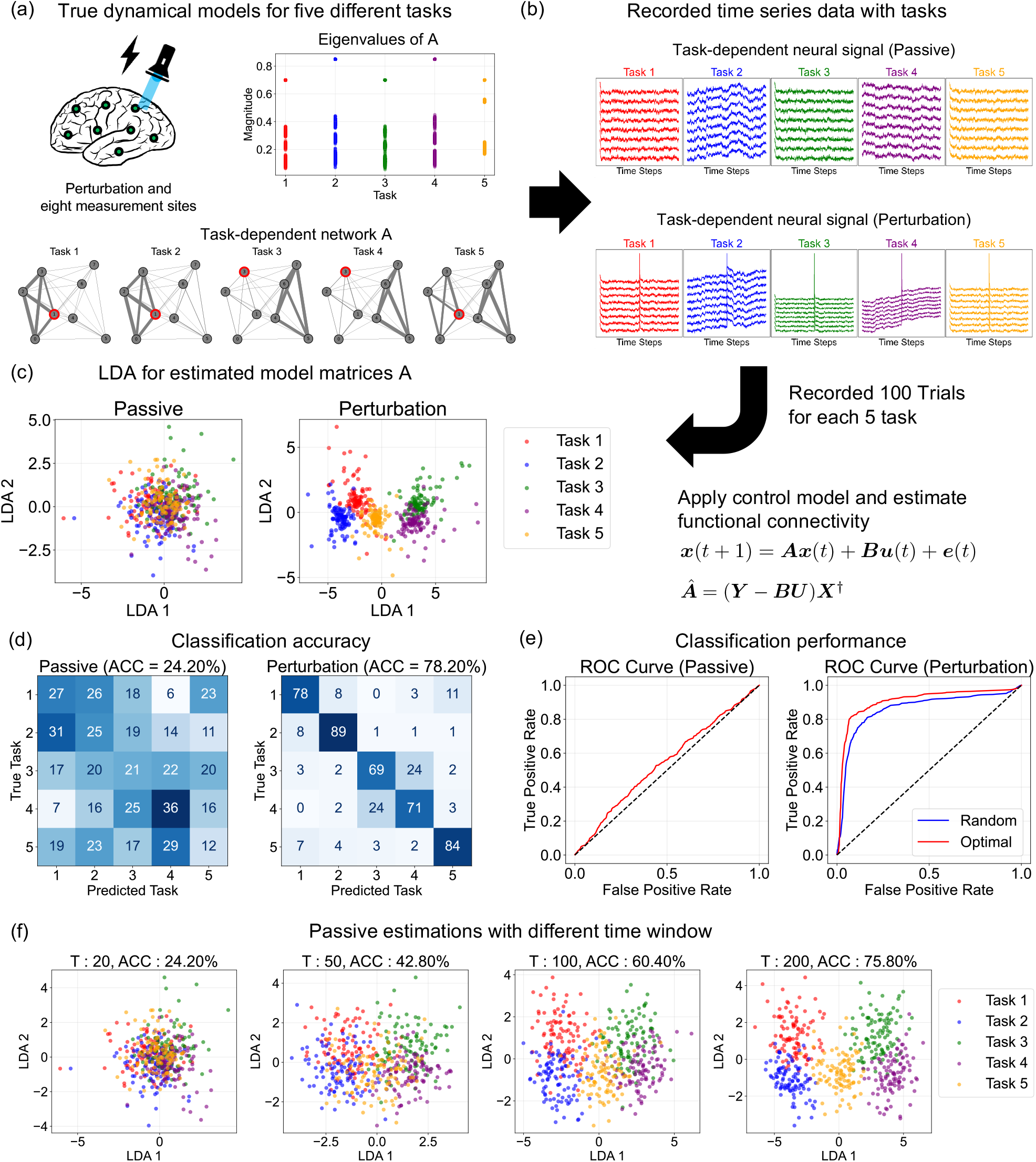
Neural state classification enhanced by perturbation-aided model estimation. (a) Ground-truth dynamics models defining five distinct task conditions, each with unique eigenvalue distributions and network structures of matrix ***A***. The optimally excited node is highlighted in red. (b) Simulated neural states (activity) induced by the five task conditions, shown separately for the passive (top) and perturbed (bottom) regimes. (c) Estimated model matrices ***A*** visualized using LDA. (d) Confusion matrices showing classification accuracy for the passive condition (left) and the optimally designed perturbation condition (right). (e) ROC curves for the passive condition (left) and a comparison of random and optimal perturbation conditions (right). (f) LDA plots for passive condition with increasing time window length, showing clustering and classification accuracy as the time window length *T* increases from 20 to 200.

The simulation demonstrates that classification performance under perturbation is superior to that under the passive condition, both in visual inspection and in quantitative evaluation. As revealed by linear discriminant analysis (LDA) projection, task clusters in the passive condition exhibit substantial overlap, whereas perturbation leads to clear separation among the tasks (Fig. 7c). Classification with LDA and 10-fold cross-validation shows the average accuracy is substantially higher in the perturbation condition (78.20%) compared to the passive condition (24.20%), as shown in Fig. 7d. ROC curve analysis further confirms that perturbation substantially improves discriminability across all tasks and outperforms the random perturbation condition (Fig. 7e). These results can be explained by Eq. 8: the designed perturbation inputs increase the eigenvalues of the covariance matrix, including even the smallest ones, which in turn leads to a decrease in the estimation error of ***A***.

To obtain estimates under the passive condition that are comparable to those derived under perturbation, it is necessary to experimentally observe extensive time-series data. Figure 7f illustrates the LDA projection and classification accuracy for different time-series lengths. The leftmost LDA plot (*T* = 20) corresponds to the passive condition shown in Fig. 7c, indicating that the estimation performance in the passive condition becomes comparable to that in the perturbation condition only when the time window reaches approximately *T* = 200, a 10-fold increase compared to *T* = 20. While the absolute duration depends on the interpretation of the time unit, such time windows may not be prohibitive in some experimental settings. Nevertheless, our results consistently show that passive observation requires substantially longer recordings to achieve comparable performance, highlighting the efficiency of the perturbation-based approach when the available data length is limited.

### Neural State Transitions via Optimal Control

This section demonstrates the practical utility of our framework by applying it to a neural state control problem [42, 51]. Here, we estimate the system parameters from both passive and perturbation states. Then, we determine optimal control inputs which control the neural states to desired targets, and associated control costs. By using perturbation inputs, ***A*** is accurately estimated, which in turn allows precise determination of the optimal inputs and the corresponding controlled state transitions (Fig. 8a). Moreover, use of the perturbation approach to derive the model allows the controllability Gramian [42, 47], as well as the estimation of control costs for each network area, to be more reliably assessed (Fig. 8b and 8c). This simulation underscores the importance of accurate system identification for achieving optimal control and reliable estimation of neural functions. Throughout this section, we use the term *perturbation input* to refer to the input used for system identification and *control input* to refer to the input used for state transitions.

**Fig. 8:**
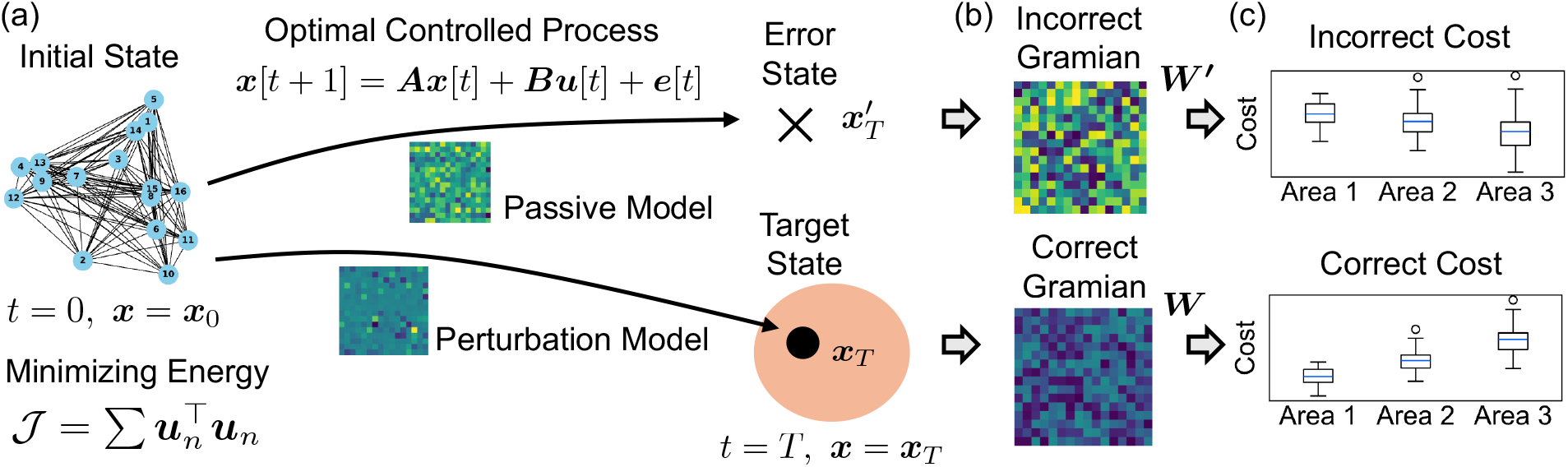
Effects of system identification on network control theory. (a) State transitions using models identified from the passive and perturbed conditions. (b) Controllability Gramians with two models. (c) Control costs computed from the estimated controllability Gramians.

We designed a network as shown in Fig. 9a to clearly show the differences in system identification between passive and perturbations states. The network consists of two disconnected groups. Nodes 1 and 2, as well as nodes 3, 4, and 5, are each connected separately, with no connections between these groups, as shown in matrix ***A***. We assumed a known input matrix ***B*** in which nodes 3 and 4 cannot be directly controlled. In this scenario, the optimal control strategy to manipulate nodes 3 and 4 is to apply inputs to node 5. However, if the system identification is inaccurate, there is the possibility that control will be attempted through nodes 1 and 2, which are disconnected from nodes 3 and 4. These matrices were estimated from the time series signals generated by simulation. We generated passive and perturbation states to estimate the parameter matrices. The perturbation condition uses an impulse input with sufficient intensity (*α* = 10^3^). By using the estimated system model, we determined optimal control inputs, then evaluated the state transitions and associated costs. The controlled transition test was run with *T* = 50. The control objective was to set nodes 3 and 4 to 25 while keeping all other nodes at 0 without any movement.

**Fig. 9:**
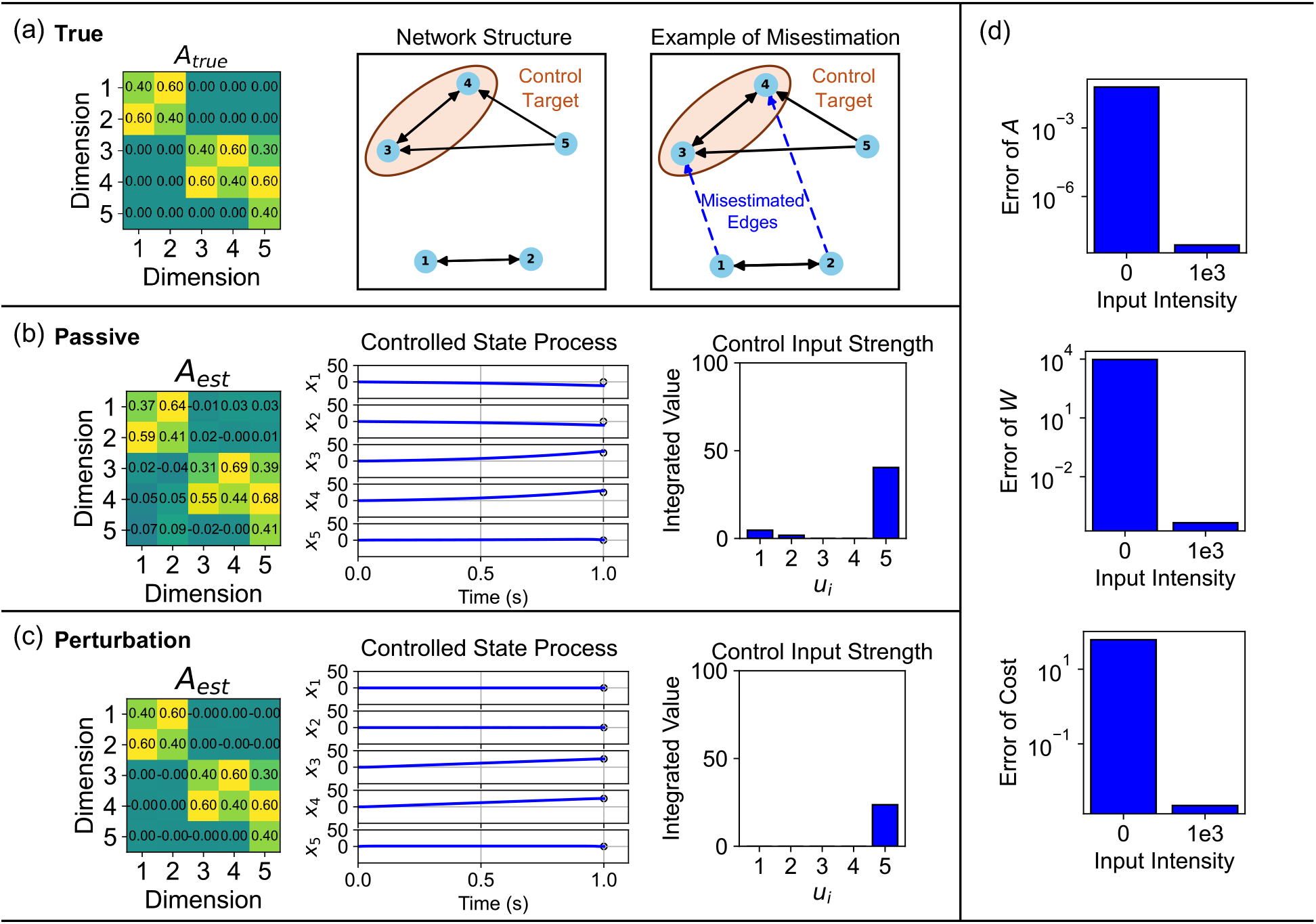
Accurate system identification via perturbation enables precise optimal control. (a) True matrix ***A*** and its network structure. Nodes 1, 2, and 5 are controllable nodes. Nodes 3 and 4 are targets controlled by the determined control input. When the connectivity matrix is misestimated, the structure has edges between nodes 1 and 2, and nodes 3 and 4. (b) Matrix estimation failure under passive condition and controlled state transitions. The control objectives are plotted as white circles. (c) Successful matrix estimations and controlled state transitions with sufficiently strong impulse inputs (*α* = 10^3^). (d) Errors in the estimated matrices ***A*** and ***W*** and the control cost (average controllability).

The simulation results show that perturbation-based estimation enables the precise control of neural states. If the estimation of ***A*** is inaccurate, the state transitions under optimal control cannot be executed correctly (Fig. 9b). Nodes 3 and 4 fail to reach the target, and unnecessary activations occur in nodes 1 and 2. Furthermore, unnecessary control inputs are required in nodes 1 and 2. On the other hand, when the model is correctly estimated using a high-strength impulse, the resulting control inputs enable accurate state transitions (Fig. 9c). In this case, the strength of the impulse is correctly concentrated on node 5. As the error in matrix ***A*** increases, the estimation errors of the controllability Gramian ***W*** lead to inaccuracies in estimating the control cost (Fig. 9d). Here, we adopt average controllability as an example measure of control cost [42, 46]. The key point here is that, since the controllability Gramian involves multiple products of ***A*** (Eq. 21), even small errors of ***A*** can result in significant discrepancies. Therefore, when estimating optimal control input for neural transitions or control costs, it is more appropriate to identify those model parameters with at least arbitrary perturbations, and ideally with designed perturbations.

### Practical Designing Procedure for Optimal Perturbation Inputs

This section presents a practical framework for designing optimal perturbation inputs to estimate neural dynamics. The practical framework progressively refines the input signal through repeated cycles of model identification and perturbation input design. By leveraging this alternating scheme, each iteration utilizes the current model estimate to design a more informative input for the next round of identification. We also focus on perturbations characterized by a broad and uniform frequency distribution, a form commonly adopted in control engineering and conceptually analogous to transcranial random noise stimulation (tRNS) in neuroscience [71, 72]. As shown in Figs. 5 and B.4, the designed composite-wave input allows for more accurate model estimation than the uniform-frequency input when their total input energies are equal.

Iterative refinement of both the perturbation design and the estimation process progressively improves the accuracy of ***A***. The time-series data is collected from 32 points, where the matrix ***A*** (Fig. 10a) is designed to have 16 oscillatory modes (i.e., 16 complex-conjugate eigenvalue pairs, yielding 32 eigenvalues in total). In this simulation, one stimulation session is applied to a single node at each design step. At each iteration, the stimulation is designed as a composite-frequency sinusoidal input encompassing all modes of the estimated ***Â***, and the target node of the perturbation is determined through numerical optimization that minimizes 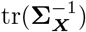. Table B.1 summarizes the simulation conditions including measurement duration and number of trials used in each simulation. The stimulation duration was set equal to the measurement duration.

**Fig. 10:**
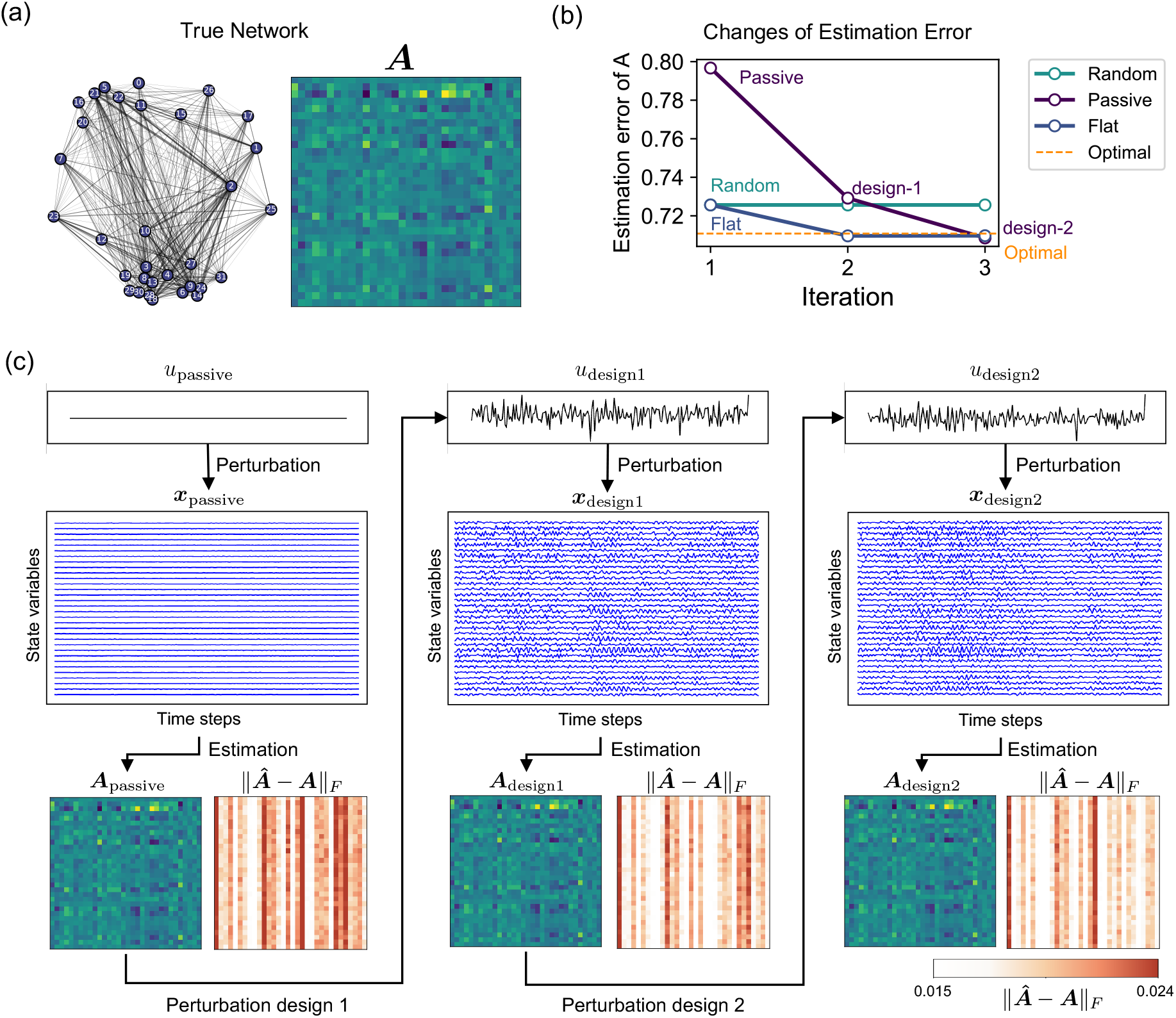
Iterative perturbation design converges to the theoretical optimum in a 32-channel neural network simulation. (a) True matrix ***A*** and its network structure, which was designed to include both oscillatory and attenuating components. (b) Comparison of estimation errors. The purple line indicates the iterative design process initialized from the passive-state estimate (Iteration 1). The blue line shows the iterative design starting from a flat-spectrum input (Iteration 1). Both processes converge towards the theoretically optimal error (dashed line) as the design is refined (design-1, design-2). The third line (labeled “Random”) represents a baseline in which a single randomly chosen node is stimulated with a flat-spectrum input throughout all iterations. (c) Schematic of the iterative procedure. An initial estimated system matrix (***A***_passive_) is used to design the first perturbation input (***u***_design1_), which yields a better estimate (***A***_design1_), and the cycle repeats.

The following procedure outlines a practical approach (Fig. 10c) for this simulation, aiming to precisely estimate the matrix by designing an optimal perturbation input (assuming frequency stimulation, such as tACS).

1. Record spontaneous activity as the passive state and estimate ***A***_passive_.
2. Based on the estimated matrix, the optimal frequency and target node of the perturbation input are determined through numerical optimization following Eq. 9.
3. The designed perturbation ***u***_design1_ is applied to the system, and a more precise matrix ***A***_design1_ is identified.
4. The obtained matrix ***A***_design1_ is again used to design a new perturbation input ***u***_design2_ and estimate a more accurate matrix ***A***_design2_. This iterative process can be continued until convergence criteria, such as changes in ***A***, are met.

The proposed framework improved estimation accuracy iteration by iteration. As shown in Fig. 10b, the estimation error converges toward the theoretical optimum (dashed line) with each design iteration. Here, the theoretical optimum is obtained from a simulation in which the input is constructed as a composite sinusoid covering all dynamical modes of the true matrix ***A***, and the stimulation location is numerically selected to minimize 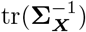. The error from the second iteration (2nd design) is smaller than that from the first (1st design), which in turn is substantially lower than the initial estimate. This convergence holds both when starting from the passive condition and when starting from a flat-spectrum input (a practical approach analogous to tRNS). While the flat-spectrum input (blue line, Iteration 1) provided a better starting point than passive data, the designed input outperformed it by the second iteration. Furthermore, the converged estimation error was lower than that of a random stimulation baseline—in which a single randomly chosen node was stimulated using a flat-spectrum (all-frequency) input throughout all iterations (green line)—whose estimation error remained approximately constant across iterations. This contrast directly demonstrates that the benefit of the proposed framework arises specifically from model-based optimization of both the spatial (node selection) and spectral (frequency composition) stimulation parameters. These results should be interpreted as a case-specific illustration rather than a universal gain, as the magnitude of improvement depends on factors such as network structure, recording duration, noise level, and stimulation design. This simulation demonstrates that our perturbation design framework enables the step-by-step refinement of system identification even without prior knowledge of the system.

## Discussion

This study addressed the challenge of designing optimal perturbations for effectively identifying neural system dynamics. We introduced a framework for estimating the optimal perturbation input for identification of neural systems. Our findings demonstrated that incorporating perturbation inputs, including TMS, tDCS, and tACS, significantly improves identification accuracy. Specifically, alignment of their parameters with the intrinsic frequencies of matrix ***A***, high input intensity, and targeted inputs directed at nodes that efficiently excite heavily damped dynamical modes (either nodes directly associated with those modes or hub nodes whose outgoing connections propagate the perturbation throughout the network) enhances system identification. Furthermore, we outlined an approach for designing the optimal perturbation input by iterating system identifications with perturbations, and demonstrated that the approach progressively decreases parameter estimation error. Beyond the parameter identification of neural systems, our findings provide insights into how these estimated parameters influence optimal control theory in neuroscience. These results emphasize the potential of perturbation input design to advance understanding of neural dynamics.

The assumption of linearity and first-order autoregressiveness in neural dynamics warrants discussion. Biological signals often exhibit complex and nonlinear interactions that cannot be fully captured by linear models. Nevertheless, prior studies have demonstrated that nonlinear behaviors can frequently be approximated using linear models [9, 73, 42]. Linear models are particularly effective for providing accurate approximations of nonlinear systems within a specific operating range [74]. Therefore, establishing theoretical foundations based on linear assumptions can contribute to the development of theories for nonlinear systems or be effectively utilized in their advancement. To extend these theories into the nonlinear domain, it would be necessary to incorporate advanced frameworks such as the Koopman operator [75, 76] and system identification methods leveraging machine learning techniques [77, 78]. On the other hand, discussions regarding higher-order VAR models are relatively straightforward due to the simplicity of the estimation method. Previous studies involving actual functional data have employed VAR models of an order greater than one [79, 80, 81]. The framework proposed in this paper can be naturally and meaningfully extended to higher-order VAR models, broadening its applicability to more complex temporal dependencies.

Safety and experimental constraints play a pivotal role in the practical implementation of this framework, influencing both the scope and design of potential applications. In biological systems, stimulation parameters, such as amplitude, frequency, and location, must adhere to stringent safety thresholds to avoid adverse effects, such as tissue damage or unintended physiological responses [82, 17]. For stimulus intensity, our proposed framework suggested that a stronger stimulus improves system identification; however, an excessively strong stimulus poses safety risks [83]. With respect to sinusoidal input, stimuli are typically applied within ranges corresponding to neural activity. The optimal input frequencies derived from the proposed theory are expected to align with these ranges, suggesting that the proposed framework can be implemented without major complications [83, 84]. Furthermore, the selection of stimulation sites is often dictated by experimental accessibility or ethical considerations [85, 86]. Based on these constraints, it is necessary to determine the most appropriate stimulation parameters to achieve optimal system identification.

Two directions warrant further investigation: extending the framework to partial observability, and validating it through stimulation experiments. In experimental neuroscience, recordings are often limited to a subset of neural populations, resulting in partial observability. A growing body of work has leveraged delay-embedding techniques, represented by Takens’ embedding theorem [87], to reconstruct hidden dynamics from partial observations [88, 89, 90, 91]. Applying such techniques enables the estimation of the full connectivity matrix, thereby extending our framework to settings with partial observability. The second direction concerns experimental validation. Validating a theoretical framework through experimental design is an essential in bridging the gap between theory and practice. Recent research involving some of the present authors has demonstrated that TMS can facilitate the discrimination of neural states [47]. Future experiments can build on this finding by incorporating passive, arbitrary, and designed optimal stimuli to estimate the connectivity matrix, and then comparing the ability of these matrices to distinguish different neural states. If these proposed evaluations demonstrate the practical effectiveness of our theory, it could then be applied to actual experiments. As shown in Fig. 10, a preliminary connectivity matrix is obtained through preliminary experiments to design optimal perturbations. These perturbations would then be applied in the main experiments, improving the estimation of connectivity matrices and neural dynamics. Experiments using our approach will enable the more precise and comprehensive analysis of brain and neural function, and in turn facilitate valuable new insights into human cognition and behavior.

## Methods

### State Vector Simulation and Model Estimation

To investigate appropriate perturbation inputs for the identification of neural systems, we constructed neural dynamics using matrices ***A*** and ***B*** to generate the temporal evolution of state variables. From the generated dynamics, we estimated the model matrix ***A***, while assuming that the model matrix ***B*** was known.

Given an initial condition ***x***(0), the neural dynamics were generated according to the true matrices ***A*** and ***B*** and the governing equation for ***x***(*t*) (Eq. 1), with a predefined noise time series ***ξ***(*t*) added at each time step. For the discrete-time simulations, the sampling rate was set to an appropriate value for each simulation. The model matrix ***A*** was then estimated from the generated time-series states ***x***(*t*) and applied perturbation inputs ***u***(*t*) using the OLS.

Both the generation of neural dynamics and the model estimation were repeated for a predefined number of trials, with variations in initial conditions and noise sequences. The estimation errors of the matrices were quantified using the Frobenius norm. Details of the simulation parameters are provided in the Supplementary Material B.1.

### Eigenvector Alignment for eigenvalue-scaled ellipsoids

To examine the relationship between the eigenvalues *µ*_*i*_ of the state covariance matrix **Σ**_***X***_ and the input frequencies, we plotted an eigenvalue-scaled ellipsoid, thereby providing an intuitive representation of optimal input design. However, the eigenvectors of **Σ**_***X***_ are not fixed; they may vary depending on the applied input, which complicates direct comparison across conditions. To enable consistent comparison of eigenvalues across input conditions, we reordered eigenvalues according to the similarity of their associated eigenvectors to those obtained in the passive condition, which served as the reference basis. For each reference eigenvector ***t***_*i*_, we identified the most similar eigenvector ***u***_*j*_ from the input condition by maximizing the absolute inner product

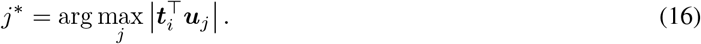

The eigenvalue *µ*_*j*\*_ corresponding to ***u***_*j*\*_ was then assigned to the (i)-th position of the reordered list. Each ***u***_*j*_ was used only once, ensuring a one-to-one correspondence between reference and input eigenvectors. This alignment resolves the permutation and sign ambiguities inherent in eigendecomposition, and allows eigenvalues to be compared across conditions along a common axis.

### Optimal Control and Network Controllability

Accurately estimating dynamical models under perturbation not only facilitates precise inference of the neural dynamics but also contributes to the accurate estimation of control inputs for neural state transitions. Recently, the control theory which utilizes the controllability Gramian and control costs is applied for neuroscience, and provides insights into the efficiency and feasibility of inducing specific neural state transitions [42, 51] with some limitations [92, 93]. For clarity, we refer to the input for system identification as the *perturbation input* and the input for state control in the optimal control theory as the *control input*.

The optimal control input is derived under the condition of assuming the linear system is controllable. We consider the sequence of control inputs that minimizes the following cost function (input energy minimization) while driving the state from the initial state ***x***_0_ to the terminal state ***x***_*T*_:

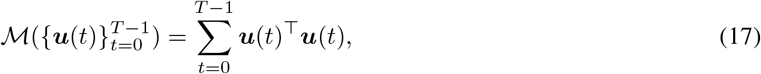

under

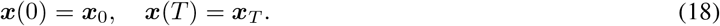

Here, the initial and terminal state conditions represent constraints on the optimization problem. This problem is expressed as follows:

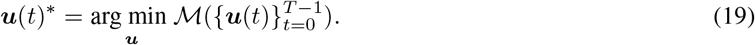

The constrained optimization problem can be solved using Lagrange multipliers. The optimal control input is given by

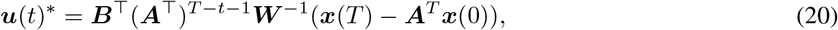

where ***W*** is the controllability Gramian, defined as:

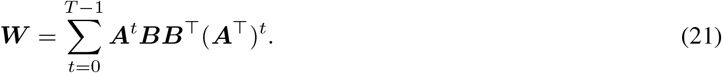

The controllability Gramian plays a pivotal role in determining the optimal control and control input cost. It has been used to identify the functional roles of individual brain regions [42, 47].

### Average Controllability for Control Cost

While the optimal control problem (Eq. 19) calculates the specific input energy for a given state transition, a more general, state-independent metric is often used to characterize the system’s overall controllability. This metric, average controllability, is a state-independent metric used to quantify the system’s overall ease of control [42, 46]. It is defined as the trace of the controllability Gramian:

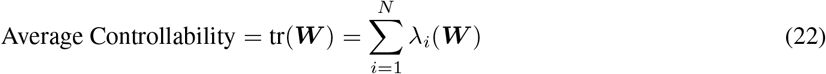

A larger trace indicates that the system is, on average, more controllable (i.e., can be moved to various states with less input energy). This metric is therefore used as a measure of control efficiency.

## Acknowledgments

We are grateful to Yumi Shikauchi, Shunsuke Kamiya, and Daiki Kiyooka for their insightful feedback and valuable discussions, which significantly contributed to the development of this research. This work was supported by JST Moonshot R&D Grant Number JPMJMS2012, JSPS KAKENHI Grant Number 24K20462 and JSPS KAKENHI Grant Number 23KJ0799.

## References

[1] Waites, A. B., Briellmann, R. S., Saling, M. M., Abbott, D. F. & Jackson, G. D. Functional connectivity networks are disrupted in left temporal lobe epilepsy. Ann. Neurol. 59, 335–343 (2006).

[2] Friston, K. J. Functional and effective connectivity: a review. Brain Connect. 1, 13–36 (2011).

[3] Boly, M. et al. Brain connectivity in disorders of consciousness. Brain Connect. 2, 1–10 (2012).

[4] Sporns, O. Contributions and challenges for network models in cognitive neuroscience. Nat. Neurosci. 17, 652–660 (2014).

[5] Petersen, S. E. & Sporns, O. Brain networks and cognitive architectures. Neuron 88, 207–219 (2015).

[6] Sporns, O. Networks of the brain. The MIT Press (MIT Press, London, England, 2016).

[7] Seguin, C. et al. Communication dynamics in the human connectome shape the cortex-wide propagation of direct electrical stimulation. Neuron 111, 1391–1401.e5 (2023).

[8] Stevens, M. C. The developmental cognitive neuroscience of functional connectivity. Brain Cogn. 70, 1–12 (2009).

[9] Honey, C. J. et al. Predicting human resting-state functional connectivity from structural connectivity. Proc. Natl. Acad. Sci. U. S. A. 106, 2035–2040 (2009).

[10] Rubinov, M. & Sporns, O. Complex network measures of brain connectivity: uses and interpretations. Neuroimage 52, 1059–1069 (2010).

[11] Friston, K., Moran, R. & Seth, A. K. Analysing connectivity with granger causality and dynamic causal modelling. Curr. Opin. Neurobiol. 23, 172–178 (2013).

[12] Siegle, J. et al. Survey of spiking in the mouse visual system reveals functional hierarchy. Nature 592, 86–92 (2021).

[13] Yan, Y. & Murphy, T. H. Decoding state-dependent cortical-cerebellar cellular functional connectivity in the mouse brain. Cell Rep. 43, 114348 (2024).

[14] Park, A. H. et al. Optogenetic mapping of functional connectivity in freely moving mice via insertable wrapping electrode array beneath the skull. ACS Nano 10, 2791–2802 (2016).

[15] Kucyi, A. et al. Intracranial electrophysiology reveals reproducible intrinsic functional connectivity within human brain networks. J. Neurosci. 38, 4230–4242 (2018).

[16] Greicius, M. D. et al. Resting-state functional connectivity in major depression: abnormally increased contributions from subgenual cingulate cortex and thalamus. Biol. Psychiatry 62, 429–437 (2007).

[17] Rocchi, F. et al. Increased fMRI connectivity upon chemogenetic inhibition of the mouse prefrontal cortex. Nat. Commun. 13, 1056 (2022).

[18] Nentwich, M. et al. Functional connectivity of EEG is subject-specific, associated with phenotype, and different from fMRI. Neuroimage 218, 117001 (2020).

[19] Bogéa Ribeiro, L. & da Silva Filho, M. Systematic review on EEG analysis to diagnose and treat autism by evaluating functional connectivity and spectral power. Neuropsychiatr. Dis. Treat. 19, 415–424 (2023).

[20] Reid, A. T. et al. Advancing functional connectivity research from association to causation. Nat. Neurosci. 22, 1751–1760 (2019).

[21] Goebel, R., Roebroeck, A.Kim, D.-S. & Formisano, E. Investigating directed cortical interactions in time-resolved fMRI data using vector autoregressive modeling and granger causality mapping. Magn. Reson. Imaging 21, 1251–1261 (2003).

[22] Ding, M., Chen, Y. & Bressler, S. L. Handbook of time series analysis: Recent theoretical developments and applications (Wiley-VCH Verlag, Weinheim, Germany, 2006), 1 edn.

[23] Ting, C., Seghouane, A., Khalid, M. U. & Salleh, S. Is first-order vector autoregressive model optimal for fMRI data? Neural Comput. 27, 1857–1871 (2015).

[24] Seth, A., Barrett, A. & Barnett, L. Granger causality analysis in neuroscience and neuroimaging. J. Neurosci. 35, 3293–3297 (2015).

[25] Cekic, S., Grandjean, D. & Renaud, O. Time, frequency, and time-varying granger-causality measures in neuroscience. Stat. Med. 37, 1910–1931 (2018).

[26] Ruiz, S., Buyukturkoglu, K., Rana, M., Birbaumer, N. & Sitaram, R. Real-time fMRI brain computer interfaces: self-regulation of single brain regions to networks. Biol. Psychol. 95, 4–20 (2014).

[27] Bassett, D. & Sporns, O. Network neuroscience. Nat. Neurosci. 20, 353–364 (2017).

[28] Lee, M.Yoon, J.-G. & Lee, S.-W. Predicting motor imagery performance from resting-state EEG using dynamic causal modeling. Front. Hum. Neurosci. 14, 321 (2020).

[29] Bergmann, T. O. et al. Concurrent TMS-fMRI for causal network perturbation and proof of target engagement. Neuroimage 237, 118093 (2021).

[30] Siddiqi, S. H., Kording, K. P., Parvizi, J. & Fox, M. D. Causal mapping of human brain function. Nature reviews neuroscience 23, 361–375 (2022).

[31] Casali, A. G. et al. A theoretically based index of consciousness independent of sensory processing and behavior. Sci. Transl. Med. 5 (2013).

[32] Stepaniants, G., Brunton, B. W. & Kutz, J. N. Inferring causal networks of dynamical systems through transient dynamics and perturbation. Phys. Rev. E. 102, 042309 (2020).

[33] Lepperød, M. E., Stöber, T., Hafting, T., Fyhn, M. & Kording, K. P. Inferring causal connectivity from pairwise recordings and optogenetics. PLoS Comput. Biol. 19, e1011574 (2023).

[34] Steinberg, E. E. & Janak, P. H. Establishing causality for dopamine in neural function and behavior with optogenetics. Brain Res. 1511, 46–64 (2013).

[35] Lepperød, M. E., Stöber, T., Hafting, T., Fyhn, M. & Kording, K. P. Inferring causal connectivity from pairwise recordings and optogenetics. PLoS Comput. Biol. 19, e1011574 (2023).

[36] Hallett, M. et al. Contribution of transcranial magnetic stimulation to assessment of brain connectivity and networks. Clin. Neurophysiol. 128, 2125–2139 (2017).

[37] Tik, M. et al. Acute TMS/fMRI response explains offline TMS network effects - an interleaved TMS-fMRI study. Neuroimage 267, 119833 (2023).

[38] Massimini, M. et al. Triggering sleep slow waves by transcranial magnetic stimulation. Proc. Natl. Acad. Sci. U. S. A. 104, 8496–8501 (2007).

[39] George, M. S. Whither TMS: A one-trick pony or the beginning of a neuroscientific revolution? Am. J. Psychiatry 176, 904–910 (2019).

[40] Bernal-Casas, D., Lee, H. J., Weitz, A. J. & Lee, J. H. Studying brain circuit function with dynamic causal modeling for optogenetic fMRI. Neuron 93, 522–532.e5 (2017).

[41] Grimm, C. et al. Tonic and burst-like locus coeruleus stimulation distinctly shift network activity across the cortical hierarchy. Nat. Neurosci. 27, 2167–2177 (2024).

[42] Gu, S. et al. Controllability of structural brain networks. Nat. Commun. 6, 8414 (2015).

[43] Deng, S., Li, J., Thomas Yeo, B. T. & Gu, S. Control theory illustrates the energy efficiency in the dynamic reconfiguration of functional connectivity. Commun. Biol. 5, 295 (2022).

[44] Kawakita, G., Kamiya, S., Sasai, S., Kitazono, J. & Oizumi, M. Quantifying brain state transition cost via schrödinger bridge. Netw. Neurosci. 6, 118–134 (2022).

[45] Kamiya, S., Kawakita, G., Sasai, S., Kitazono, J. & Oizumi, M. Optimal control costs of brain state transitions in linear stochastic systems. J. Neurosci. 43, 270–281 (2023).

[46] Moradi Amani, A. et al. Controllability of functional and structural brain networks. Complexity 2024 (2024).

[47] Shikauchi, Y. et al. Quantifying state-dependent control properties of brain dynamics from perturbation responses. J. Neurosci. e0364252025 (2025).

[48] Minai, Y., Smith, M., Soldado-Magraner, J. & Yu, B. MiSO: Optimizing brain stimulation to create neural activity states. In Globerson, A.et al. (eds.) Advances in Neural Information Processing Systems 37, vol. 37, 24126–24149 (Neural Information Processing Systems Foundation, Inc. (NeurIPS), San Diego, California, USA, 2024).

[49] Wagenmaker, A. et al. Active learning of neural population dynamics using two-photon holographic optogenetics. Adv. Neural Inf. Process. Syst. 37, 31659–31687 (2024).

[50] Angulo-Sherman, I. N., Rodríguez-Ugarte, M., Sciacca, N., Iáñez, E. & Azorín, J. M. Effect of tDCS stimulation of motor cortex and cerebellum on EEG classification of motor imagery and sensorimotor band power. J. Neuroeng. Rehabil. 14, 31 (2017).

[51] Karrer, T. M. et al. A practical guide to methodological considerations in the controllability of structural brain networks. J. Neural Eng. 17, 026031 (2020).

[52] Betzel, R. F., Gu, S., Medaglia, J. D., Pasqualetti, F. & Bassett, D. S. Optimally controlling the human connectome: the role of network topology. Sci. Rep. 6, 30770 (2016).

[53] Kim, J. Z. et al. Role of graph architecture in controlling dynamical networks with applications to neural systems. Nat. Phys. 14, 91–98 (2018).

[54] Braun, U. et al. Brain network dynamics during working memory are modulated by dopamine and diminished in schizophrenia. Nat. Commun. 12, 3478 (2021).

[55] Ahmed, S. & Nozari, E. On the linearizing effect of spatial averaging in large-scale populations of homogeneous nonlinear systems. In 2022 IEEE 61st Conference on Decision and Control (CDC) (IEEE, 2022).

[56] Nozari, E. et al. Macroscopic resting-state brain dynamics are best described by linear models. Nat. Biomed. Eng. 8, 68–84 (2024).

[57] Momi, D., Wang, Z. & Griffiths, J. D. TMS-evoked responses are driven by recurrent large-scale network dynamics. Elife 12 (2023).

[58] Tapsell, L. C. et al. What are the optimal transcranial direct current stimulation parameters and design elements to modulate corticospinal excitability? a systematic review and longitudinal meta-analysis. Neurol. Res. Pract. 7, 86 (2025).

[59] Grover, S., Fayzullina, R., Bullard, B. M., Levina, V. & Reinhart, R. M. G. A meta-analysis suggests that tACS improves cognition in healthy, aging, and psychiatric populations. Sci. Transl. Med. 15, eabo2044 (2023).

[60] Ogata, K. Modern Control Engineering (Prentice Hall, 2010).

[61] Nise, N. S. Control Systems Engineering (John Wiley & Sons, 2020), 8 edn.

[62] Lozano, A. M. et al. Deep brain stimulation: current challenges and future directions. Nat. Rev. Neurol. 15, 148–160 (2019).

[63] Deisseroth, K. Optogenetics. Nat. Methods 8, 26–29 (2011).

[64] Tong, L. et al. Single cell in vivo optogenetic stimulation by two-photon excitation fluorescence transfer. iScience 26, 107857 (2023).

[65] LaFosse, P. K. et al. Single-cell optogenetics reveals attenuation-by-suppression in visual cortical neurons. bioRxivorg 2023.09.13.557650 (2024).

[66] Green, M. & Moore, J. B. Persistence of excitation in linear systems. Syst. Control Lett. 7, 351–360 (1986).

[67] Shimkin, N. & Feuer, A. Persistency of excitation in continuous-time systems. Syst. Control Lett. 9, 225–233 (1987).

[68] Jenkins, B. M., Annaswamy, A. M., Lavretsky, E. & Gibson, T. E. Convergence properties of adaptive systems and the definition of exponential stability. SIAM J. Control Optim. 56, 2463–2484 (2018).

[69] Lu, X. & Cannon, M. Robust adaptive model predictive control with persistent excitation conditions. Automatica (Oxf.) 152, 110959 (2023).

[70] Hamilton, J. D. Time Series Analysis (Princeton University Press, Princeton, 1994).

[71] Paulus, W. Transcranial electrical stimulation (tES - tDCS; tRNS, tACS) methods. Neuropsychol. Rehabil. 21, 602–617 (2011).

[72] Paulus, W., Nitsche, M. A. & Antal, A. Application of transcranial electric stimulation (tDCS, tACS, tRNS): From motor-evoked potentials towards modulation of behaviour. Eur. Psychol. 21, 4–14 (2016).

[73] Fernández Galán, R. On how network architecture determines the dominant patterns of spontaneous neural activity. PLoS One 3, e2148 (2008).

[74] Khalil, H. K. Nonlinear systems (Prentice-Hall, Upper Saddle River, NJ, 2002).

[75] Koopman, B. O. Hamiltonian systems and transformation in hilbert space. Proc. Natl. Acad. Sci. U. S. A. 17, 315–318 (1931).

[76] Chow, C., Dan, T., Styner, M. & Wu, G. Understanding brain dynamics through neural koopman operator with structure-function coupling: 27th international conference, marrakesh, morocco, october 6–10, 2024, proceedings, part II. In Linguraru, M. G.et al. (eds.) Medical Image Computing and Computer Assisted Intervention – MICCAI 2024, vol. 15002 of Lecture Notes in Computer Science, 509–518 (Springer Nature Switzerland, Cham, 2024).

[77] Suk, H.-I., Wee, C.-Y.Lee, S.-W. & Shen, D. State-space model with deep learning for functional dynamics estimation in resting-state fMRI. Neuroimage 129, 292–307 (2016).

[78] Glaser, J. I., Benjamin, A. S., Farhoodi, R. & Kording, K. P. The roles of supervised machine learning in systems neuroscience. Prog. Neurobiol. 175, 126–137 (2019).

[79] Tseng, S. Y., Chen, R. C., Chong, F. C. & Kuo, T. S. Evaluation of parametric methods in EEG signal analysis. Med. Eng. Phys. 17, 71–78 (1995).

[80] Chang, J.-Y. et al. Multivariate autoregressive models with exogenous inputs for intracerebral responses to direct electrical stimulation of the human brain. Front. Hum. Neurosci. 6, 317 (2012).

[81] Shakeel, A., Onojima, T., Tanaka, T. & Kitajo, K. Real-time implementation of EEG oscillatory phase-informed visual stimulation using a least mean square-based AR model. J. Pers. Med. 11, 38 (2021).

[82] Bikson, M. et al. Safety of transcranial direct current stimulation: Evidence based update 2016. Brain Stimul. 9, 641–661 (2016).

[83] Antal, A. et al. Low intensity transcranial electric stimulation: Safety, ethical, legal regulatory and application guidelines. Clin. Neurophysiol. 128, 1774–1809 (2017).

[84] Mager, T. et al. High frequency neural spiking and auditory signaling by ultrafast red-shifted optogenetics. Nat. Commun. 9, 1750 (2018).

[85] Rossi, S., Hallett, M., Rossini, P. M., Pascual-Leone, A. & Safety of TMS Consensus Group. Safety, ethical considerations, and application guidelines for the use of transcranial magnetic stimulation in clinical practice and research. Clin. Neurophysiol. 120, 2008–2039 (2009).

[86] Heinrichs, J.-H. The promises and perils of non-invasive brain stimulation. Int. J. Law Psychiatry 35, 121–129 (2012).

[87] Takens, F. Detecting strange attractors in turbulence. In Rand, D. & Young, L.-S. (eds.) Dynamical Systems and Turbulence, Warwick 1980, vol. 898 of Lecture Notes in Mathematics, 366–381 (Springer, Berlin, Heidelberg, 1981).

[88] Arbabi, H. & Mezić, I. Computation of transient koopman spectrum using hankel-dynamic mode decompoisition. APS G1.009 (2017).

[89] Brunton, S. L., Brunton, B. W., Proctor, J. L., Kaiser, E. & Kutz, J. N. Chaos as an intermittently forced linear system. Nat. Commun. 8, 19 (2017).

[90] Ostrow, M., Eisen, A. & Fiete, I. Delay embedding theory of neural sequence models. arXiv [cs.LG] (2024).

[91] Huang, A. et al. InputDSA: Demixing then comparing recurrent and externally driven dynamics. arXiv [q-bio.NC] (2025).

[92] Tu, C. et al. Warnings and caveats in brain controllability. Neuroimage 176, 83–91 (2018).

[93] Suweis, S. et al. Brain controllability: Not a slam dunk yet. Neuroimage 200, 552–555 (2019).

