## Supplementary Material for "Designing optimal perturbation inputs for system identification in neuroscience"

### A Theoretical Details

#### A.1 Asymptotic Property of the Estimator for $A$

This section derives the asymptotic estimation error of the OLS estimator $\hat{\boldsymbol{A}}$ for the system matrix $\boldsymbol{A}$. The primary goal is to obtain a first-order asymptotic approximation for the Mean Squared Error (MSE) of the estimation, quantified by the expected squared Frobenius norm, $\mathbb{E}[\|\hat{\boldsymbol{A}} - \boldsymbol{A}\|_F^2]$.

We consider the linear dynamical system

$$\boldsymbol{x}(t+1) = \boldsymbol{A}\boldsymbol{x}(t) + \boldsymbol{B}\boldsymbol{u}(t) + \boldsymbol{\xi}(t), \tag{S.1}$$

where the noise $\boldsymbol{\xi}(t)$ is i.i.d. over time with zero mean and covariance matrix $\boldsymbol{\Sigma}_\xi = \mathbb{E}[\boldsymbol{\xi}(t)\boldsymbol{\xi}(t)^\top] \in \mathbb{R}^{n\times n} \succ 0$.

We estimate $\boldsymbol{A}$ by OLS from the data matrices $\boldsymbol{X} \in \mathbb{R}^{n\times T}$, $\tilde{\boldsymbol{Y}} := \boldsymbol{Y} - \boldsymbol{B}\boldsymbol{U} \in \mathbb{R}^{n\times T}$, giving $\hat{\boldsymbol{A}} = \tilde{\boldsymbol{Y}}\boldsymbol{X}^\dagger$.

The asymptotic distribution of the vectorized estimation error of the OLS estimator $\hat{\boldsymbol{A}}$ is given by [1], Proposition 11.1 and Appendix 11.A:

$$\sqrt{T}\big(\mathrm{vec}(\hat{\boldsymbol{A}}) - \mathrm{vec}(\boldsymbol{A})\big) \xrightarrow{L} \mathcal{N}\big(\boldsymbol{0},\, \boldsymbol{\Sigma}_{\mathrm{Asym}}\big), \tag{S.2}$$

where $T$ is the number of samples, $\xrightarrow{L}$ denotes convergence in distribution, and $\boldsymbol{\Sigma}_{\mathrm{Asym}}$ is the asymptotic covariance matrix. Following the notation in the original text (and its assumptions), this covariance is:

$$\boldsymbol{\Sigma}_{\mathrm{Asym}} = \boldsymbol{\Sigma}_\xi \otimes \boldsymbol{\Sigma}_{\boldsymbol{X}}^{-1}, \tag{S.3}$$

where $\otimes$ denotes the Kronecker product [1, Appendix A.4, p. 732], $\boldsymbol{\Sigma}_\xi \in \mathbb{R}^{n\times n}$ is the noise covariance, and $\boldsymbol{\Sigma}_{\boldsymbol{X}} = \mathbb{E}[\boldsymbol{x}(t)\boldsymbol{x}(t)^\top] \in \mathbb{R}^{n\times n}$ is the state covariance matrix, so that $\boldsymbol{\Sigma}_{\mathrm{Asym}} \in \mathbb{R}^{n^2\times n^2}$.

We are interested in the estimation error measured by the squared Frobenius norm, $\|\hat{\boldsymbol{A}} - \boldsymbol{A}\|_F^2$. Let us define the normalized error vector $\boldsymbol{z}_T$ based on (S.2):

$$\boldsymbol{z}_T := \sqrt{T}\big(\mathrm{vec}(\hat{\boldsymbol{A}}) - \mathrm{vec}(\boldsymbol{A})\big). \tag{S.4}$$

We know that $\boldsymbol{z}_T \xrightarrow{L} \boldsymbol{z}$, where $\boldsymbol{z} \sim \mathcal{N}(\boldsymbol{0}, \boldsymbol{\Sigma}_{\mathrm{Asym}})$ is an $n^2$-dimensional Gaussian vector.

The squared Frobenius norm can be expressed as the squared Euclidean norm of the vectorized error:

$$\|\hat{\boldsymbol{A}} - \boldsymbol{A}\|_F^2 = \|\mathrm{vec}(\hat{\boldsymbol{A}} - \boldsymbol{A})\|_2^2. \tag{S.5}$$

Multiplying by $T$, we can relate this quantity to $\boldsymbol{z}_T$:

$$T \cdot \|\hat{\boldsymbol{A}} - \boldsymbol{A}\|_F^2 = \Big(\sqrt{T} \cdot \|\mathrm{vec}(\hat{\boldsymbol{A}} - \boldsymbol{A})\|_2\Big)^2 = \boldsymbol{z}_T^\top \boldsymbol{z}_T. \tag{S.6}$$

By the continuous mapping theorem, since $g(\boldsymbol{z}) = \boldsymbol{z}^\top\boldsymbol{z}$ is a continuous function, the convergence in distribution of $\boldsymbol{z}_T$ implies the convergence in distribution of $\boldsymbol{z}_T^\top\boldsymbol{z}_T$:

$$T \cdot \|\hat{\boldsymbol{A}} - \boldsymbol{A}\|_F^2 \xrightarrow{L} \boldsymbol{z}^\top\boldsymbol{z}, \quad \text{where} \quad \boldsymbol{z} \sim \mathcal{N}(\boldsymbol{0}, \boldsymbol{\Sigma}_{\mathrm{Asym}}). \tag{S.7}$$

Its limit of the expected value equals the expectation of the limiting distribution $\boldsymbol{z}^\top\boldsymbol{z}$:

$$\lim_{T\to\infty} \mathbb{E}\Big[T \cdot \|\hat{\boldsymbol{A}} - \boldsymbol{A}\|_F^2\Big] = \mathbb{E}\big[\boldsymbol{z}^\top\boldsymbol{z}\big] \tag{S.8}$$

$$= \mathbb{E}\big[\mathrm{tr}(\boldsymbol{z}\boldsymbol{z}^\top)\big] \tag{S.9}$$

$$= \mathrm{tr}\big(\mathbb{E}[\boldsymbol{z}\boldsymbol{z}^\top]\big)\,. \tag{S.10}$$

By definition, $\mathbb{E}[\boldsymbol{z}\boldsymbol{z}^\top]$ is the covariance matrix of the limiting distribution $\boldsymbol{z}$, which is $\boldsymbol{\Sigma}_{\mathrm{Asym}}$.

$$\lim_{T\to\infty} \mathbb{E}\Big[T \cdot \|\hat{\boldsymbol{A}} - \boldsymbol{A}\|_F^2\Big] = \mathrm{tr}(\boldsymbol{\Sigma}_{\mathrm{Asym}}) \tag{S.11}$$

$$= \mathrm{tr}(\boldsymbol{\Sigma}_\xi \otimes \boldsymbol{\Sigma}_{\boldsymbol{X}}^{-1}). \tag{S.12}$$

Using the trace property $\mathrm{tr}(\boldsymbol{A}\otimes\boldsymbol{B}) = \mathrm{tr}(\boldsymbol{A})\,\mathrm{tr}(\boldsymbol{B})$, we obtain:

$$\lim_{T\to\infty} \mathbb{E}\Big[T \cdot \|\hat{\boldsymbol{A}} - \boldsymbol{A}\|_F^2\Big] = \mathrm{tr}(\boldsymbol{\Sigma}_\xi)\,\mathrm{tr}(\boldsymbol{\Sigma}_{\boldsymbol{X}}^{-1}). \tag{S.13}$$

This result implies that for a sufficiently large $T$, the expected estimation error can be approximated as:

$$\mathbb{E}\left[\|\hat{\boldsymbol{A}} - \boldsymbol{A}\|_F^2\right] \approx \frac{1}{T}\operatorname{tr}(\boldsymbol{\Sigma}_\xi)\operatorname{tr}(\boldsymbol{\Sigma}_{\boldsymbol{X}}^{-1}). \tag{S.14}$$

This expression provides a simple interpretation of the estimation error. It confirms that the error is inversely proportional to the sample size $T$, meaning the error decreases as more data is collected. The error is also directly proportional to the total noise power, $\operatorname{tr}(\boldsymbol{\Sigma}_\xi)$, indicating that more noise leads to more error. Furthermore, the error is proportional to $\operatorname{tr}(\boldsymbol{\Sigma}_{\boldsymbol{X}}^{-1})$, which acts as an inverse measure of signal strength; a weaker signal $\boldsymbol{x}(t)$ (i.e., less variation) leads to a larger error. This relationship resembles the inverse of the Signal-to-Noise ratio, where the error is governed by the balance between noise, signal strength, and the amount of data.

#### A.2 State Vectors Considering Typical Inputs Used in Neuroscience

##### A.2.1 Sinusoidal Input

To determine which frequencies most effectively enhance system identification accuracy, we now analyze the time evolution of the state vector $\boldsymbol{x}(t)$ in response to a sinusoidal inputs. This analysis is the necessary first step toward the goal established in the main text (i.e., sufficiently exciting the system's dynamics to increase the eigenvalues of the covariance matrix). By understanding how $\boldsymbol{x}(t)$ itself behaves under different frequencies, we can identify which inputs are most effective. We therefore examine frequency external inputs, represented in neuroscience by tACS, to derive the optimal input frequency based on the system's dynamic response.

In the case where the perturbation consists of a composite sinusoidal input comprising multiple frequencies, the input can be written as

$$\boldsymbol{u}(k) = \sum_{l=1}^{L}\cos(\omega_l k)\boldsymbol{u}_0, \tag{S.15}$$

where $\omega_k$ is the angular frequencies.

The general solution which we denote as Eq. 10 is:

$$\boldsymbol{x}(t) = \boldsymbol{A}^t\boldsymbol{x}(0) + \sum_{k=0}^{t-1}\boldsymbol{A}^{t-1-k}\boldsymbol{B}\boldsymbol{u}(k) + \sum_{k=0}^{t-1}\boldsymbol{A}^{t-1-k}\boldsymbol{\xi}(k) \tag{S.16}$$

Inserting Eq. S.15 into Eq. 10 yields:

$$\boldsymbol{x}(t) = \boldsymbol{A}^t\boldsymbol{x}(0) + \sum_{k=0}^{t-1}\boldsymbol{A}^{t-1-k}\boldsymbol{B}\sum_{l=1}^{L}\cos(\omega_l k)\boldsymbol{u}_0 + \sum_{k=0}^{t-1}\boldsymbol{A}^{t-1-k}\boldsymbol{\xi}(k) \tag{S.17}$$

Under the assumption that the system is stable (all eigenvalues $\lambda_d$ of $\boldsymbol{A}$ satisfy $|\lambda_d| < 1$), the second term of Eq. S.17 is the critical component to analyze.

By performing diagonalization of $\boldsymbol{A}$ as $\boldsymbol{A} = \boldsymbol{V}\boldsymbol{\Lambda}\boldsymbol{V}^{-1}$, where $\boldsymbol{V}$ is the matrix of eigenvectors, $\boldsymbol{\Lambda}$ is the diagonal matrix of eigenvalues $\lambda_d$, ... The spectral decomposition of $\boldsymbol{A}^m$ is $\boldsymbol{A}^m = \sum_{d=1}^{n}\lambda_d^m\boldsymbol{v}_d\boldsymbol{w}_d^\top$, where $\boldsymbol{v}_d$ denotes the $d$-th column vector of $\boldsymbol{V}$, and $\boldsymbol{w}_d^\top$ denotes the $d$-th row vector of $\boldsymbol{V}^{-1}$. The second term of Eq. S.17 can be expressed as follows:

$$\sum_{k=0}^{t-1}\boldsymbol{A}^{\,t-1-k}\boldsymbol{B}\sum_{l=1}^{L}\cos(\omega_l k)\boldsymbol{u}_0 \tag{S.18}$$

$$= \sum_{k=0}^{t-1}\boldsymbol{V}\boldsymbol{\Lambda}^{\,t-1-k}\boldsymbol{V}^{-1}\boldsymbol{B}\sum_{l=1}^{L}\cos(\omega_l k)\boldsymbol{u}_0 \tag{S.19}$$

$$= \sum_{k=0}^{t-1}\sum_{d=1}^{n}\lambda_d^{\,t-1-k}\,\boldsymbol{v}_d\boldsymbol{w}_d^\top\boldsymbol{B}\sum_{l=1}^{L}\cos(\omega_l k)\boldsymbol{u}_0 \tag{S.20}$$

Using Euler's formula ($\cos x = (e^{ix} + e^{-ix})/2$) and swapping the order of summation (which is permissible by linearity), we get:

$$\sum_{k=0}^{t-1} \sum_{d=1}^{n} \lambda_d^{t-1-k} \, \boldsymbol{v}_d \boldsymbol{w}_d^\top \boldsymbol{B} \sum_{l=1}^{L} \cos(\omega_l k) \boldsymbol{u}_0 \tag{S.21}$$

$$= \sum_{k=0}^{t-1} \sum_{d=1}^{n} \sum_{l=1}^{L} \frac{e^{i\omega_l k} + e^{-i\omega_l k}}{2} \, \lambda_d^{t-1-k} \, \boldsymbol{v}_d \boldsymbol{w}_d^\top \boldsymbol{B} \, \boldsymbol{u}_0 \tag{S.22}$$

$$= \frac{1}{2} \sum_{l=1}^{L} \sum_{d=1}^{n} \left( \sum_{k=0}^{t-1} \lambda_d^{t-1-k} e^{i\omega_l k} + \sum_{k=0}^{t-1} \lambda_d^{t-1-k} e^{-i\omega_l k} \right) \boldsymbol{v}_d \boldsymbol{w}_d^\top \boldsymbol{B} \, \boldsymbol{u}_0 \tag{S.23}$$

We now introduce the polar form of the eigenvalues, $\lambda_d = r_d e^{i\theta_d}$. Given the stability assumption, we have $r_d = |\lambda_d| < 1$. This yields:

$$= \frac{1}{2} \sum_{l=1}^{L} \sum_{d=1}^{n} \left( \frac{\lambda_d^t - e^{i\omega_l t}}{\lambda_d - e^{i\omega_l}} + \frac{\lambda_d^t - e^{-i\omega_l t}}{\lambda_d - e^{-i\omega_l}} \right) \boldsymbol{v}_d \boldsymbol{w}_d^\top \boldsymbol{B} \, \boldsymbol{u}_0 \tag{S.24}$$

$$= \frac{1}{2} \sum_{l=1}^{L} \sum_{d=1}^{n} \left( \frac{r_d^t e^{i\theta_d t} - e^{i\omega_l t}}{r_d e^{i\theta_d} - e^{i\omega_l}} + \frac{r_d^t e^{i\theta_d t} - e^{-i\omega_l t}}{r_d e^{i\theta_d} - e^{-i\omega_l}} \right) \boldsymbol{v}_d \boldsymbol{w}_d^\top \boldsymbol{B} \, \boldsymbol{u}_0 \tag{S.25}$$

$$= \frac{1}{2} \sum_{l=1}^{L} \sum_{d=1}^{n} \left( \frac{(r_d^t e^{i\theta_d t} - e^{i\omega_l t})(r_d e^{-i\theta_d} - e^{-i\omega_l})}{(r_d e^{i\theta_d} - e^{i\omega_l})(r_d e^{-i\theta_d} - e^{-i\omega_l})} + \frac{(r_d^t e^{i\theta_d t} - e^{-i\omega_l t})(r_d e^{-i\theta_d} - e^{i\omega_l})}{(r_d e^{i\theta_d} - e^{-i\omega_l})(r_d e^{-i\theta_d} - e^{i\omega_l})} \right) \boldsymbol{v}_d \boldsymbol{w}_d^\top \boldsymbol{B} \, \boldsymbol{u}_0 \tag{S.26}$$

$$= \frac{1}{2} \sum_{l=1}^{L} \sum_{d=1}^{n} \left( \frac{r_d^{t+1} e^{i\theta_d (t-1)} - r_d^t e^{i(\theta_d t - \omega_l)} - r_d e^{i(\omega_l t - \theta_d)} + e^{i\omega_l (t-1)}}{r_d^2 - 2 r_d \cos(\theta_d - \omega_l) + 1} \right.$$
$$\left. + \frac{r_d^{t+1} e^{i\theta_d (t-1)} - r_d^t e^{i(\theta_d t + \omega_l)} - r_d e^{i(-\omega_l t - \theta_d)} + e^{-i\omega_l (t-1)}}{r_d^2 - 2 r_d \cos(\theta_d + \omega_l) + 1} \right) \boldsymbol{v}_d \boldsymbol{w}_d^\top \boldsymbol{B} \, \boldsymbol{u}_0, \tag{S.27}$$

For brevity, we define the following terms:

$$Q(\omega_l, \theta_d) = \frac{r_d^{t+1} e^{i\theta_d (t-1)} - r_d^t e^{i(\theta_d t - \omega_l)} - r_d e^{i(\omega_l t - \theta_d)} + e^{i\omega_l (t-1)}}{r_d^2 - 2 r_d \cos(\theta_d - \omega_l) + 1} \tag{S.28}$$

$$\boldsymbol{F}_d = \boldsymbol{v}_d \boldsymbol{w}_d^\top \boldsymbol{B} \boldsymbol{u}_0 \tag{S.29}$$

The second term is:

$$\sum_{k=0}^{t-1} \boldsymbol{A}^{t-1-k} \boldsymbol{B} \sum_{l=1}^{L} \cos(\omega_l k) \boldsymbol{u}_0 = \frac{1}{2} \sum_{l=1}^{L} \sum_{d=1}^{n} \left( Q(\omega_l, \theta_d) + Q(-\omega_l, \theta_d) \right) \boldsymbol{F}_d \tag{S.30}$$

(S.31)

Since the matrix $\boldsymbol{A}$ is real, its complex eigenvalues and eigenvectors must appear in complex conjugate pairs. If $\lambda_d = r_d e^{i\theta_d}$ (with $\theta_d \neq 0$) is an eigenvalue, then $\lambda_{d'} = \overline{\lambda_d} = r_d e^{-i\theta_d}$ is also an eigenvalue for some index $d'$. Thus, $\theta_{d'} = -\theta_d$. Furthermore, since $\boldsymbol{B}$ and $\boldsymbol{u}_0$ are also real, the vector $\boldsymbol{F}_d = \boldsymbol{v}_d \boldsymbol{w}_d^\top \boldsymbol{B} \boldsymbol{u}_0$ associated with $\lambda_d$ and the vector $\boldsymbol{F}_{d'} = \boldsymbol{v}_{d'} \boldsymbol{w}_{d'}^\top \boldsymbol{B} \boldsymbol{u}_0$ associated with $\lambda_{d'}$ are also complex conjugates: $\boldsymbol{F}_{d'} = \overline{\boldsymbol{F}_d}$. We can use this property to simplify the sum.

$$Q(-\omega_l, \theta_{d'}) \boldsymbol{F}_{d'} = Q(-\omega_l, -\theta_d) \overline{\boldsymbol{F}_d} \tag{S.32}$$

$$= \frac{r_d^{t+1} e^{-i\theta_d (t-1)} - r_d^t e^{i(-\theta_d t + \omega_l)} - r_d e^{i(-\omega_l t + \theta_d)} + e^{-i\omega_l (t-1)}}{r_d^2 - 2 r_d \cos(-\theta_d + \omega_l) + 1} \overline{\boldsymbol{F}_d} \tag{S.33}$$

$$= \overline{\frac{r_d^{t+1} e^{i\theta_d (t-1)} - r_d^t e^{i(\theta_d t - \omega_l)} - r_d e^{i(\omega_l t - \theta_d)} + e^{i\omega_l (t-1)}}{r_d^2 - 2 r_d \cos(\theta_d - \omega_l) + 1} \boldsymbol{F}_d} \tag{S.34}$$

$$= \overline{Q(\omega_l, \theta_d) \boldsymbol{F}_d} \tag{S.35}$$

Similarly, $Q(-\omega_l, \theta_d)\boldsymbol{F}_d = \overline{Q(\omega_l, \theta_{d'})\boldsymbol{F}_{d'}}$

For real eigenvalues ($\theta_d = 0$), $\boldsymbol{F}_d$ is real, and we find:

$$Q(-\omega_l, 0)\boldsymbol{F}_d = \frac{r_d^{t+1} - r_d^t e^{i\omega_l} - r_d e^{-i\omega_l t} + e^{-i\omega_l (t-1)}}{r_d^2 - 2r_d \cos(\omega_l) + 1}\boldsymbol{F}_d \tag{S.36}$$

$$= \overline{\frac{r_d^{t+1} - r_d^t e^{-i\omega_l} - r_d e^{i\omega_l t} + e^{i\omega_l (t-1)}}{r_d^2 - 2r_d \cos(\omega_l) + 1}\boldsymbol{F}_d} \tag{S.37}$$

$$= \overline{Q(\omega_l, 0)\boldsymbol{F}_d} \tag{S.38}$$

We divide the set of indices of eigenvalues into three subsets as follows.Let $D_R$ denote the set of indices corresponding to real eigenvalues ($\lambda_d \in \mathbb{R}$). Among complex eigenvalues, let $D_C^+$ denote the set of indices whose imaginary part is positive, and let $D_C^-$ denote the set of indices whose imaginary part is negative.

$$\frac{1}{2}\sum_{d=1}^{n} \left(Q(\omega_l, \theta_d) + Q(-\omega_l, \theta_d)\right)\boldsymbol{F}_d = \frac{1}{2}\sum_{d \in D_R} \left(Q(\omega_l, 0)\boldsymbol{F}_d + \overline{Q(\omega_l, 0)\boldsymbol{F}_d}\right) \tag{S.39}$$

$$+ \frac{1}{2}\sum_{d \in D_C^+} \left(Q(\omega_l, \theta_d)\boldsymbol{F}_d + Q(-\omega_l, \theta_d)\boldsymbol{F}_d\right) \tag{S.40}$$

$$+ \frac{1}{2}\sum_{d' \in D_C^-} \left(Q(\omega_l, \theta_{d'})\boldsymbol{F}_{d'} + Q(-\omega_l, \theta_{d'})\boldsymbol{F}_{d'}\right) \tag{S.41}$$

Now, we pair the terms from $D_C^+$ and $D_C^-$. For each $d \in D_C^+$, there is a corresponding $d' \in D_C^-$ such that $\theta_{d'} = -\theta_d$ and $\boldsymbol{F}_{d'} = \overline{\boldsymbol{F}_d}$. Using the properties $Q(-\omega_l, \theta_d)\boldsymbol{F}_d = \overline{Q(\omega_l, \theta_{d'})\boldsymbol{F}_{d'}}$ and $Q(-\omega_l, \theta_{d'})\boldsymbol{F}_{d'} = \overline{Q(\omega_l, \theta_d)\boldsymbol{F}_d}$, the sum of the $D_C^+$ and $D_C^-$ terms becomes:

$$\frac{1}{2}\sum_{d \in D_C^+} \left(Q(\omega_l, \theta_d)\boldsymbol{F}_d + \overline{Q(\omega_l, \theta d')\boldsymbol{F}_{d'}} + Q(\omega_l, \theta d')\boldsymbol{F}_{d'} + \overline{Q(\omega_l, \theta_d)\boldsymbol{F}_d}\right) \tag{S.42}$$

$$= \frac{1}{2}\sum_{d \in D_C^+} \left((Q_d\boldsymbol{F}_d + \overline{Q_d\boldsymbol{F}_d}) + (Qd'\boldsymbol{F}_{d'} + \overline{Qd'\boldsymbol{F}_{d'}})\right) \tag{S.43}$$

$$= \sum_{d \in D_C^+} \mathrm{Re}\left[Q(\omega_l, \theta_d)\boldsymbol{F}_d\right] + \sum_{d' \in D_C^-} \mathrm{Re}\left[Q(\omega_l, \theta_{d'})\boldsymbol{F}_{d'}\right] \tag{S.44}$$

Combining this with the $D_R$ sum, which is $\sum_{d \in D_R} \mathrm{Re}[Q(\omega_l, 0)\boldsymbol{F}_d]$, we arrive at:

$$\sum_{d=1}^{n} \mathrm{Re}\left[Q(\omega_l, \theta_d)\boldsymbol{F}_d\right] \tag{S.45}$$

This is the real part of the product of the terms $Q(\omega_l, \theta_d)$ and $\boldsymbol{F}_d$ defined in Eqs. S.28 and S.29. Therefore, the second term of the $\boldsymbol{x}(t)$ can be expressed as:

$$\sum_{k=0}^{t-1} \boldsymbol{A}^{t-1-k}\boldsymbol{B}\sum_{l=1}^{L} \cos(\omega_l k)\boldsymbol{u}_0 \tag{S.46}$$

$$= \sum_{l=1}^{L}\sum_{d=1}^{n} \mathrm{Re}\left[\frac{r_d^{t+1} e^{i\theta_d (t-1)} - r_d^t e^{i(\theta_d t - \omega_l)} - r_d e^{i(\omega_l t - \theta_d)} + e^{i\omega_l (t-1)}}{r_d^2 - 2r_d \cos(\theta_d - \omega_l) + 1}\boldsymbol{v}_d\boldsymbol{w}_d^\top\boldsymbol{B}\,\boldsymbol{u}_0\right], \tag{S.47}$$

Therefore, the state vector $\boldsymbol{x}(t)$ is decomposed into a passive and a difference term as follows,

$$\boldsymbol{x}(t) = \boldsymbol{x}_{\text{passive}}(t) + \boldsymbol{x}_{\text{diff}}(t), \tag{S.48}$$

$$\boldsymbol{x}_{\text{passive}}(t) = \boldsymbol{A}^t\boldsymbol{x}(0) + \sum_{k=0}^{t-1} \boldsymbol{A}^{t-1-k}\boldsymbol{\xi}(k) \tag{S.49}$$

$$\boldsymbol{x}_{\text{diff}}(t) = \sum_{l=1}^{L}\sum_{d=1}^{n} \mathrm{Re}\left[\frac{r_d^{t+1} e^{i\theta_d (t-1)} - r_d^t e^{i(\theta_d t - \omega_l)} - r_d e^{i(\omega_l t - \theta_d)} + e^{i\omega_l (t-1)}}{r_d^2 - 2r_d \cos(\theta_d - \omega_l) + 1}\boldsymbol{v}_d\boldsymbol{w}_d^\top\boldsymbol{B}\,\boldsymbol{u}_0\right] \tag{S.50}$$

where $\boldsymbol{x}_{\text{passive}}(t)$ represents the state $\boldsymbol{x}$ that would be realized if no perturbation were applied, i.e., the state that would evolve according to Eq. 10 with $\boldsymbol{u} = 0$ in the main text. $\boldsymbol{x}_{\text{diff}}(t)$ represents the change in the state trajectory induced by the specific perturbation input $\boldsymbol{u}(k) = \sum_{l=1}^{L} \cos(\omega_l k)\boldsymbol{u}_0$. As defined in the previous section, $\lambda_d = r_d e^{i\theta_d}$ is the polar form of the $d$-th eigenvalue, the vector $\boldsymbol{v}_d$ denotes the right eigenvector of $\boldsymbol{A}$ associated with $\lambda_d$, and $\boldsymbol{w}_d^\top$ denotes the corresponding left eigenvector. The numerator of the fraction in Eq. S.47 consists of four distinct oscillatory terms. The resonant behavior of this expression is primarily governed by the denominator, $D(\omega_l, \theta_d) = r_d^2 - 2r_d\cos(\theta_d - \omega_l) + 1$. The gain term $1/D(\omega_l, \theta_d)$ has a single extremum (a maximum) with respect to $\omega_l$, which occurs when $\cos(\theta_d - \omega_l) = 1$, i.e., when the input frequency $\omega_l$ matches the eigenfrequency (argument) $\theta_d$ of the system.

##### A.2.2 Impulse and Step Input

Impulse and step inputs follow simple rules. Let $\alpha$ represent the scaling factor for impulse input. Then, input $\boldsymbol{u}(k)$ is defined as:

$$\boldsymbol{u}(k) = \begin{cases} \alpha\boldsymbol{u}_0 & \text{if } k = 0, \\ 0 & \text{if } k > 0. \end{cases} \tag{S.51}$$

Here, $\boldsymbol{u}_0 \in \mathbb{R}^m$ represents the fundamental input direction applied to the system, determining which input channel receives the impulse. We obtain the decomposition:

$$\boldsymbol{x}_{\text{passive}}(t) = \boldsymbol{A}^t\boldsymbol{x}(0) + \sum_{k=0}^{t-1} \boldsymbol{A}^{t-1-k}\boldsymbol{\xi}(k), \quad \boldsymbol{x}_{\text{diff}}(t) = \alpha\boldsymbol{A}^{t-1}\boldsymbol{B}\boldsymbol{u}_0. \tag{S.52}$$

For a scaled step input, $\boldsymbol{u}(k)$ is defined as

$$\boldsymbol{u}(k) = \beta\boldsymbol{u}_0, \quad \forall k \geq 0, \tag{S.53}$$

where $\beta$ is input strength. We then get the decomposition:

$$\boldsymbol{x}_{\text{passive}}(t) = \boldsymbol{A}^t\boldsymbol{x}(0) + \sum_{k=0}^{t-1} \boldsymbol{A}^{t-1-k}\boldsymbol{\xi}(k), \quad \boldsymbol{x}_{\text{diff}}(t) = \beta\sum_{k=0}^{t-1} \boldsymbol{A}^{t-1-k}\boldsymbol{B}\boldsymbol{u}_0. \tag{S.54}$$

From these solutions, we can infer that the input-driven terms ($\boldsymbol{x}_{\text{diff}}(t)$) are linear in the input strengths $\alpha$ and $\beta$. Consequently, the contribution of these inputs to the state covariance matrix (i.e., the outer product of the state vector) will scale quadratically with $\alpha^2$ and $\beta^2$. This analysis, which suggests that such inputs can increase the rank or magnitude of the covariance matrix eigenvalues, is explored further in the main text.

### B Experimental Conditions and Results

#### B.1 Simulation Details

The specific conditions used for generating neural dynamics in each section are summarized in Table B.1. These include factors such as sampling frequency, trial length, and perturbation input strength, which are crucial for designing reproducible simulations and evaluations. These parameters are chosen to illustrate the theoretical insights as clearly as possible, rather than to characterise the framework's behaviour across generic regimes.

#### B.2 Robustness to Nonlinear State Dynamics

We introduce nonlinearity into the state update by replacing the linear update with a nonlinear function of the current state,

$$\boldsymbol{x}(t{+}1) = \frac{\boldsymbol{A}}{\beta}\tanh\big(\beta\,\boldsymbol{x}(t)\big) + \boldsymbol{B}\boldsymbol{u}(t) + \boldsymbol{\xi}(t), \tag{S.1}$$

on a 4-dimensional two-mode network, which is the extended setup of Fig. 2. The detailed parameters are given in Table B.1. The $1/\beta$ prefactor normalises the dynamics. The $\tanh$ function is a common choice for introducing nonlinearity [2] while maintaining bounded outputs, and the parameter $\beta$ controls the degree of nonlinearity (Fig. B.1a). Because the true map is now nonlinear in $\boldsymbol{x}$, a direct comparison of $\hat{\boldsymbol{A}}$ against the true $\boldsymbol{A}$ is no longer well-defined; we therefore evaluate the misspecified linear OLS estimator by the relative one-step prediction RMSE and the relative free-run (rollout) RMSE.

Under both metrics, the impulse condition tracks the true dynamics more accurately than passive observation across the swept range, with the gap shrinking when $\beta$ grows large enough that the nonlinearity itself re-excites the 40 Hz

Table B.1: Conditions for generating neural dynamics

| **Section** | Sampling Frequency | Time Length | Data Length $T$ | Number of Trials | Noise | Input Strength |
|---|---|---|---|---|---|---|
| Fig. 2 (Failure of Passive Estimation) | 1000 | 10 | 10,000 | 100 | $10^{-32}$ | 1 |
| Fig. 4-5 (Sinusoidal Inputs) | 100 | 50 | 5,000 | 10,000 | $10^{-2}$ | 1 |
| Fig. 6 (Location Tuning) | 1000 | 1 | 1,000 | 200 | $10^{-2}$ | 10 |
| Fig. 7 (Neural State Classification) | 10 | 2 | 20 | 100 | $10^{-2}$ | $10^{2}$ |
| Fig. 9 (Optimal Control) | 50 | 1 | 50 | 100 | $10^{-2}$ | $10^{3}$ |
| Fig. 10 (Practical Designing Procedure) | 100 | 25 | 2,500 | 500 | $10^{-2}$ | 0.1 |
| Fig. B.1 (Impulse/Step) | 100 | 50 | 5,000 | 1 | $10^{-2}$ | 0.5–5 |
| Fig. B.2 (White Noise / Designed) | 100 | 100 | 10,000 | 1 | $10^{-2}$ | 0.1 |
| Fig. B.3 (Nonlinear Dynamics) | 1000 | 1 | 1,000 | 30 | $10^{-32}$ | 0.15–0.3 |
| Fig. B.4 ($\boldsymbol{B}$-Matrix Uncertainty) | 1000 | 1 | 1,000 | 30 | $10^{-32}$ | 0.15–0.3 |

hidden mode strongly (Fig. B.1b,c). Figure B.1d,e show the time series for the passive and impulse conditions across the range of $\beta$, illustrating that the impulse condition strongly excites the mode dynamics. Figure B.1f,g compare the true and predicted time series; under both error metrics, the impulse condition tracks the true dynamics more accurately than passive observation. Figure B.1h shows the top three principal components of the data. Even when nonlinearity is present in the dynamics, the impulse condition still re-excites the hidden mode (third PC) and yields better predictions than passive observation.

##### B.3 Joint estimation of $A$ and $B$

The naive estimator that assumes $\boldsymbol{B} = \boldsymbol{I}$ absorbs the unmodelled term $(\boldsymbol{B}_{\text{true}} - \boldsymbol{I})\boldsymbol{u}$ into $\hat{\boldsymbol{A}}$, so its error in $\boldsymbol{A}$ grows with the mismatch level $\varepsilon = \|\boldsymbol{B}_{\text{true}} - \boldsymbol{I}\|_F$. A natural remedy is to estimate $\boldsymbol{A}$ and $\boldsymbol{B}$ jointly by regressing the next state on the stacked $[\boldsymbol{X}; \boldsymbol{U}]$ block,

$$[\hat{\boldsymbol{A}},\, \hat{\boldsymbol{B}}] \;=\; \boldsymbol{X}_+\, [\boldsymbol{X}_-;\, \boldsymbol{U}]^{\dagger}, \tag{S.2}$$

which is unbiased whenever the input excites every state coordinate. We compare seven conditions on a 4-dimensional, 2-mode discrete-time system: the ground truth; passive observation; joint estimation under four perturbation patterns, with input on (i) node 1 alone, (ii) node 3 alone, (iii) nodes 1 and 3, and (iv) all four nodes; and a baseline that uses the all-nodes input but estimates $\boldsymbol{A}$ under the naive assumption $\hat{\boldsymbol{B}} = \boldsymbol{I}$. The true matrix is same setup of Fig. 2. The detailed parameters are given in Table B.1. Every stimulated trial uses 4 impulses whose amplitudes on the active channels are drawn independently from $\mathcal{N}(0, \boldsymbol{I})$ per pulse, so that the rank of the $\boldsymbol{U}$-block of the regressor equals $\min(n_{\text{active}}, 4)$. We parametrise the mismatch as $\boldsymbol{B}_{\text{true}} = \boldsymbol{I} + \varepsilon\boldsymbol{R}$ with a fixed random matrix $\boldsymbol{R}$ normalised so that $\|\boldsymbol{R}\|_F = 1$, which makes $\varepsilon$ the scalar control parameter for the sweep.

The results show that the perturbation can still improve the estimation of $\boldsymbol{A}$ even when $\boldsymbol{B}$ is unknown, provided that $\boldsymbol{A}$ and $\boldsymbol{B}$ are recovered jointly by OLS. In the joint-estimation results of Figure B.2a, only the columns of $\boldsymbol{B}$ corresponding to the stimulated channels are recovered: the $(1, 3)$ pattern fills two columns, and the all-nodes pattern recovers $\boldsymbol{B}$ to the accuracy attainable when $\boldsymbol{B}$ is known. Nevertheless, $\boldsymbol{A}$ can be estimated correctly, including its hidden dynamical mode. The third column of the same panel shows that the naive estimator under the assumption $\hat{\boldsymbol{B}} = \boldsymbol{I}$ recovers $\boldsymbol{A}$ only approximately. Figure B.2b shows the corresponding state trajectories. Figure B.2c shows $\|\boldsymbol{A} - \hat{\boldsymbol{A}}\|_F$ as $\varepsilon$ is swept on a log scale. Passive observation is pinned at the hidden-mode bias, and the naive $\hat{\boldsymbol{B}} = \boldsymbol{I}$ baseline grows with $\varepsilon$ as the unmodelled $(\boldsymbol{B}_{\text{true}} - \boldsymbol{I})\boldsymbol{u}_t$ term inflates the residual variance. Among the joint estimators, stimulating node 1 alone yields an intermediate error because node 1 is not directly coupled to the hidden mode; stimulating node 3 yields a markedly lower error because node 3 excites the hidden mode and thereby renders it identifiable; and stimulating all nodes is better still, matching the error achievable when $\boldsymbol{B}$ is known. The $(1, 3)$ pattern lies between the node 3 and all-nodes cases. The error $\|\boldsymbol{B} - \hat{\boldsymbol{B}}\|_F$ grows linearly with $\varepsilon$ for the rank-deficient stimulation patterns and the naive baseline, and matches the known-$\boldsymbol{B}$ baseline only for the all-nodes pattern under joint OLS. Although these results provide only a preliminary demonstration of the robustness of the proposed framework, joint estimation of $\boldsymbol{A}$ and $\boldsymbol{B}$ is a promising direction for future work to relax the assumption of known $\boldsymbol{B}$.

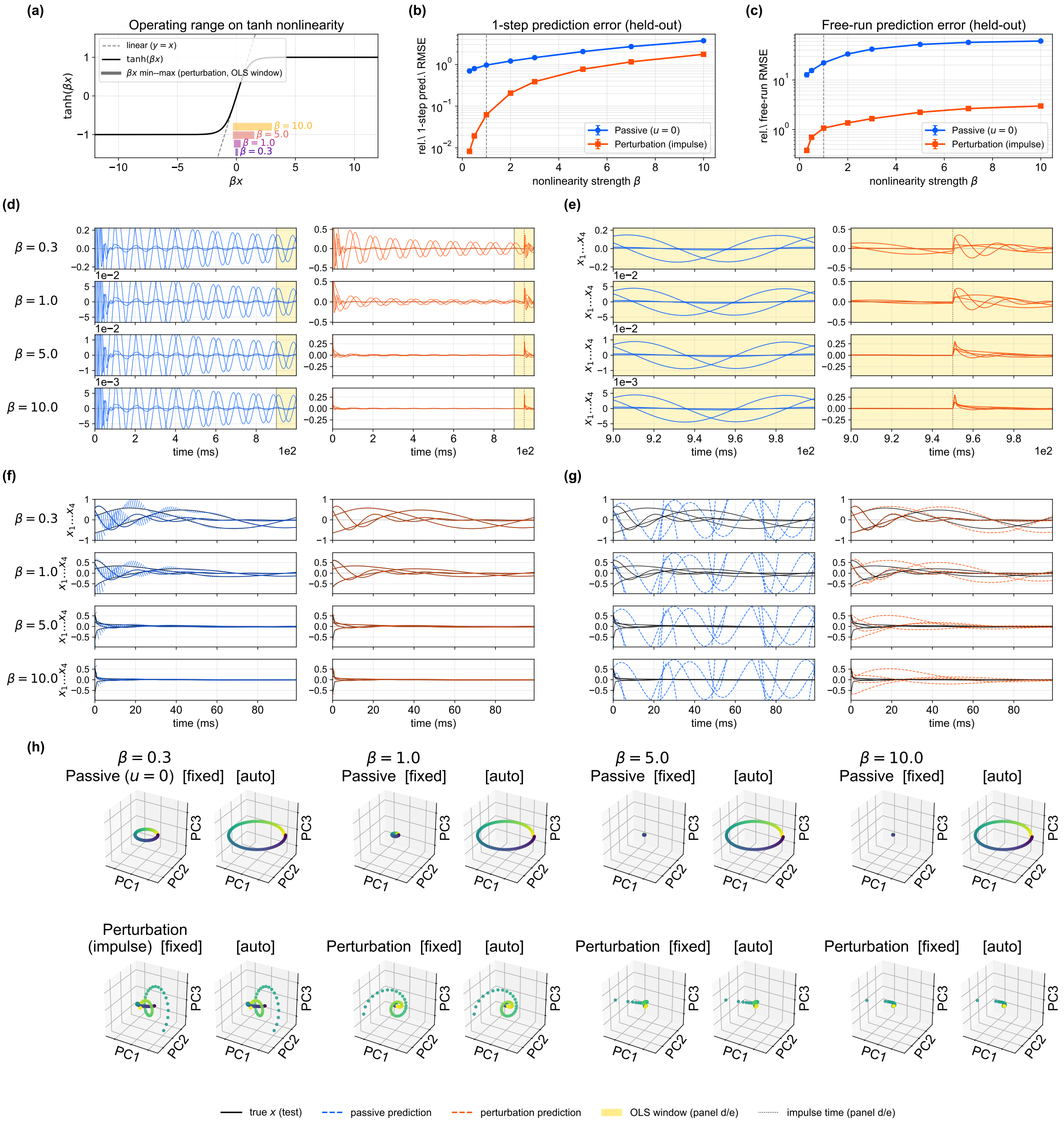

Fig. B.1: Robustness of the perturbation framework to nonlinear state dynamics. (a) The tanh nonlinearity with the min–max envelope of $\beta \boldsymbol{x}$ within the OLS window shaded for $\beta \in \{0.3, 1, 5, 10\}$. (b) Relative 1-step prediction RMSE vs. $\beta$. (c) Relative free-run (rollout) RMSE vs. $\beta$. (d) Training trajectories within the OLS estimation window for $\beta \in \{0.3, 1, 5, 10\}$, under passive observation and impulse perturbation. (e) The same trajectories shown in full; the gold shade marks the OLS window and the dotted line marks the impulse time. (f) Held-out test trajectories (black) overlaid with the per-segment 1-step prediction from $\hat{\boldsymbol{A}}, \hat{\boldsymbol{B}}$. (g) The same held-out test trajectories overlaid with the free-run (rollout) prediction. (h) The OLS-window data in the top three principal components (independent PCA per cell; viridis = sample index).

#### B.4 Demonstration of Impulse and Step Input

Impulse- and step-like inputs, such as those applied in TMS or tDCS, can substantially improve system identification accuracy when applied at high intensities. By perturbing the system state further from equilibrium, the resulting

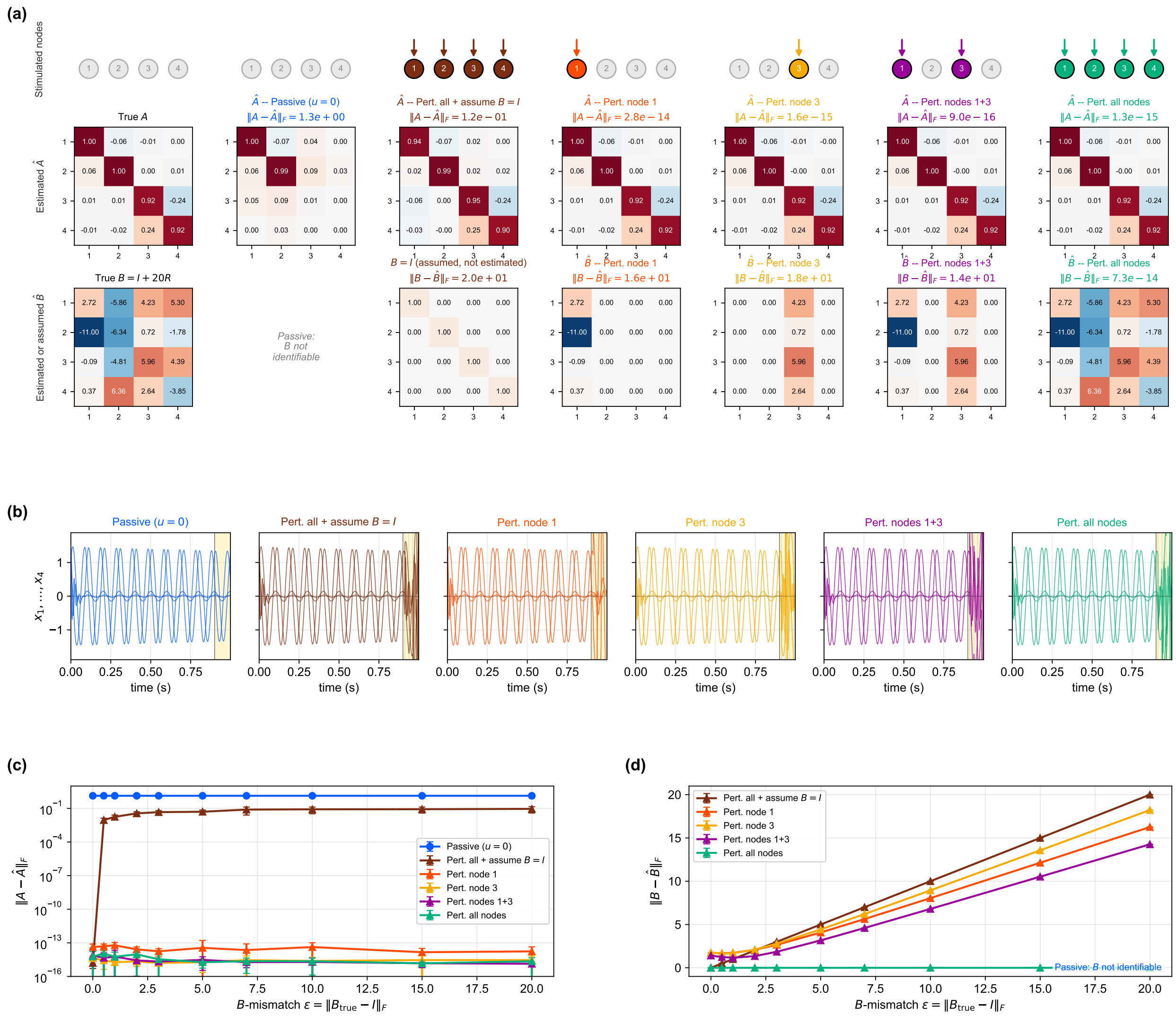

Fig. B.2: Perturbation framework can still improve the estimation of $\boldsymbol{A}$ even when $\boldsymbol{B}$ is unknown, provided that $\boldsymbol{A}$ and $\boldsymbol{B}$ are recovered jointly by OLS. (a) Top row: which channel(s) each condition stimulates (filled nodes are active; gray nodes are silent). Middle and bottom rows: recovered $\boldsymbol{A}$ and $\boldsymbol{B}$ at $\varepsilon = 20$ from a single trial. Seven conditions are compared: ground truth, passive observation, a naive baseline that uses the all-nodes input but assumes $\hat{\boldsymbol{B}} = \boldsymbol{I}$, and joint estimation under four perturbation patterns (node 1 only, node 3 only, nodes 1+3, all nodes). (b) Corresponding sample state trajectories; the gold shade marks the OLS window and dotted lines mark the impulse times. (c) $\|\boldsymbol{A} - \hat{\boldsymbol{A}}\|_F$ as a function of $\varepsilon$ (log scale). (d) $\|\boldsymbol{B} - \hat{\boldsymbol{B}}\|_F$ as a function of $\varepsilon$.

decay dynamics provide richer trajectory information. This richer information increases the eigenvalues of the state covariance matrix and, in turn, reduces estimation error.

The relationship between input intensity and system identification accuracy was examined through simulations using matrix $\boldsymbol{A}$, as illustrated in Fig. B.3a. For the dynamics generated by matrix $\boldsymbol{A}$, system identification was performed using impulse and step inputs of varying intensities. As shown in Figs. B.3b and B.3c, the identification error decreases exponentially as the input intensity increases. This error was observed to be inversely correlated with the sum of the inverses of the eigenvalues of the state covariance matrix. In practice, the strength of perturbation input must be constrained within the limits allowed by experimental conditions. However, our findings indicate that perturbation inputs should be designed to be as strong as possible within biologically acceptable ranges.

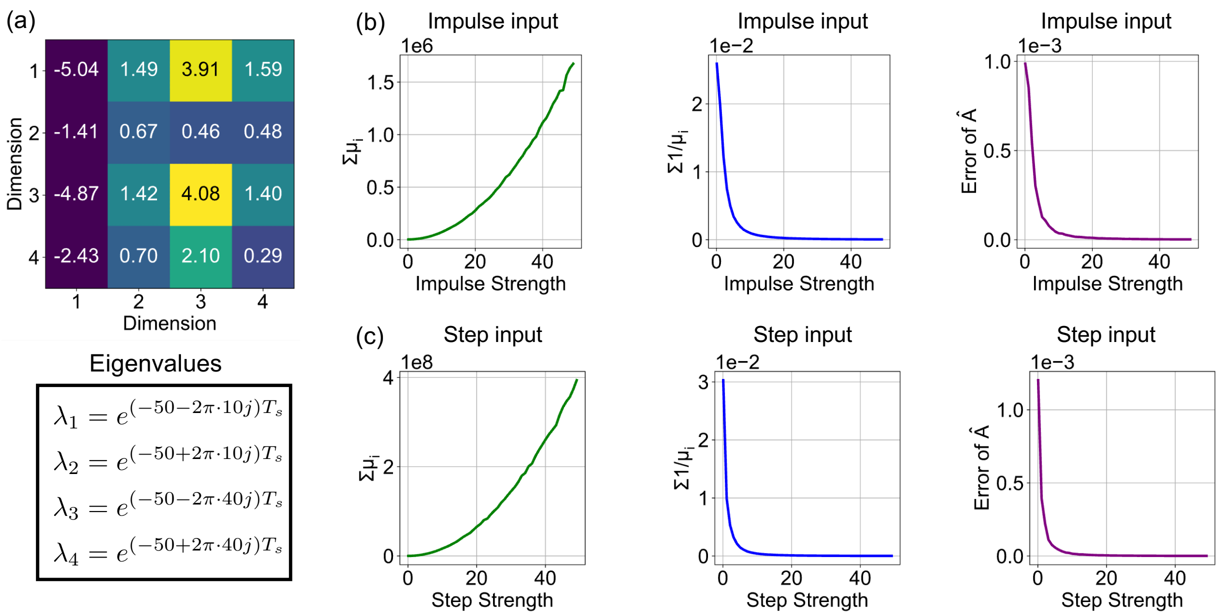

Fig. B.3: Variation of eigenvalues and system matrix error with impulse and step input strengths. (a) Assumed system matrix $\boldsymbol{A}$ and its eigenvalues. (b) Changes in the sum of eigenvalues, the sum of their reciprocals, and the estimation error of $\boldsymbol{A}$ with respect to impulse input strength. (c) Changes in the same quantities with respect to step input strength.

#### B.5 White Noise vs. Concatenated Frequency Input

We demonstrate both the effectiveness and sub-optimality of white noise input for neural system identification, a standard approach widely used in control engineering. [3, 4]. White noise contains a broad and nearly uniform range of frequencies, making it a convenient and practical choice for identifying system models when no prior information is available. However, as discussed in the main text, the input should ideally be designed to match the dynamical modes of the system. Thus, we consider white noise input as a practical yet potentially suboptimal method for neural system identification.

To assess the feasibility of the white noise perturbation input, we assume that a neural perturbation system is capable of generating random noise-like signals and that these input signals can be recorded simultaneously. We generated 32-channel time-series data using the matrix $\boldsymbol{A}$ shown in Fig. B.4a. Data were generated at sampling rate $f_s = 100$ Hz, with total length $T = 10,000$ samples. The system matrix $\boldsymbol{A} \in \mathbb{R}^{32\times 32}$ was constructed as the block diagonal of 16 discrete-time $2 \times 2$ rotation blocks whose eigenvalue radii were drawn from $\mathrm{Unif}(0.9,\ 0.95)$ and modal frequencies from $\mathrm{Unif}(1,\ 100)$ Hz. The system dynamics exhibit multiple modes, as shown in Fig. B.4b. The input matrix was $\boldsymbol{B} = \boldsymbol{I}_{32}$ with single-node stimulation applied through its first column. Process noise was i.i.d. Gaussian with per-component standard deviation $\sigma_\xi = 10^{-2}$. The designed input was a sinusoidal input with frequencies matching the modal frequencies of the system, which were extracted from the eigenvalues of $\boldsymbol{A}$. Starting from white noise, the modal frequencies $\{\omega_k\}$ extracted from the eigenvalues of $\hat{\boldsymbol{A}}$ were used as the component frequencies of the next input (design-1), and this redesign was iterated twice more (design-2, design-3). To ensure a fair comparison, both the white noise and the designed inputs were normalized to have the same total input power. This setup allows for a controlled comparison between white noise input and frequency-optimized input.

The system identification error decreased when using frequency-designed inputs rather than white noise. Figure B.4c shows the sum of reciprocal eigenvalues ($\sum_i 1/\mu_i$) of the covariance matrix of the state vectors. This value decreased when using frequency-designed inputs rather than white noise, indicating improved accuracy in system identification. Moreover, iterative redesigns progressively reduced the sum of reciprocal eigenvalues (designs 2 and 3), leading to more accurate identification. As shown in Fig. B.4d, both white noise and designed concatenated frequency inputs excited all dynamical modes without collapsing the smaller axes. The ellipsoids obtained with the designed input had smaller volumes, indicating higher identification accuracy. Therefore, this simulation demonstrates that iterative optimal design can yield more accurate system identification than conventional white noise stimulation.

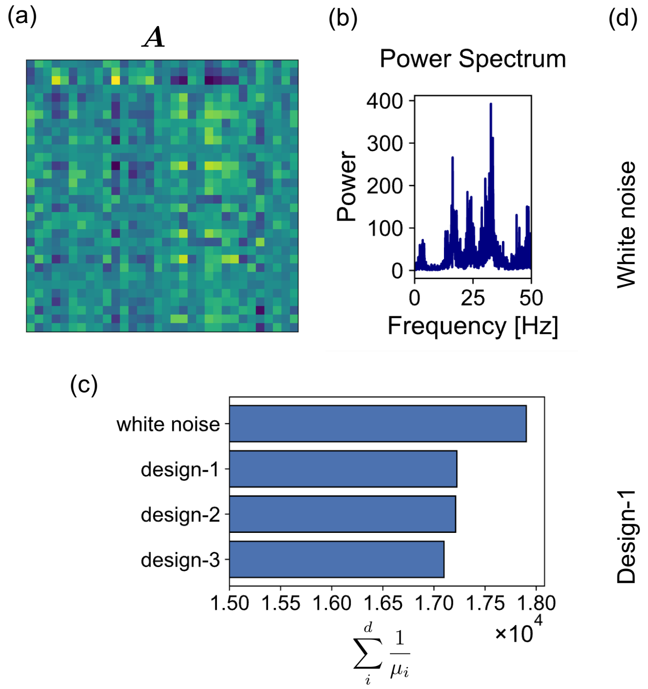

Fig. B.4: Comparison between white noise input and optimized frequency input. (a) Matrix A with 16 modes, assuming 32 channels. (b) Channel-averaged power spectrum in the passive state. (c) Comparison of the sum of reciprocal eigenvalues ($\sum_i 1/\mu_i$) of the covariance matrix of the state vector under white noise and optimized frequency input. (d) Comparison of eigenvalue-scaled ellipsoids obtained from white noise and optimized frequency input, visualizing the components corresponding to the 1st–3rd and 30th–32nd eigenvalues.

In summary, white noise input offers a simple and practical strategy for neural system identification when no prior dynamical information is available. Nevertheless, it remains suboptimal compared with concatenated frequency input, which provides higher accuracy and stability through iterative refinement. These findings also highlight the importance of input optimization in achieving precise system identification in complex neural systems.

### References

[1] Hamilton, J. D. *Time Series Analysis* (Princeton University Press, Princeton, 1994).

[2] Khalil, H. K. *Nonlinear systems* (Prentice-Hall, Upper Saddle River, NJ, 2002).

[3] Goodwin, G. C. & Payne, R. L. *Dynamic System Identification. Experiment design and Data Analysis* (Academic Press, 1977).

[4] Ljung, L. *System Identification: Theory for the User, 2nd ed* (Prentice Hall PTR, 1999).
